# Under the cover of darkness: a transcriptomic exploration of clubroot during the night

**DOI:** 10.1101/2025.01.31.635897

**Authors:** Andrea Garvetto, Susann Auer, Freia Benade, Michaela Hittorf, Jutta Ludwig-Müller, Sigrid Neuhauser

**Affiliations:** Department of Microbiology, Universität Innsbruck, Technikerstraße 25, 6020 Innsbruck, Tirol, Austria; Faculty of Biology, Technische Universität Dresden, 01062 Dresden, Germany; Department of Botany, Universität Innsbruck, Sternwartestraße 15, 6020 Innsbruck, Tirol, Austria

**Keywords:** Biotrophic parasite, Circadian, Host-pathogen interactions, *Plasmodiophora brassicae*, Protist, Time-resolved transcriptome, *Arabidopsis thaliana*

## Abstract

*Plasmodiophora brassicae* (Phytomyxea, Rhizaria) is the etiological agent of clubroot disease, one of the most important diseases of Brassicaceae crops. Alteration of metabolism and hormone homeostasis leads to the formation of tumour-like galls in the roots of affected plants. Host plant energy metabolism, defence and developmental processes are under strong temporal control and the very same processes are affected by clubroot. For the first time, this study uses time-resolved transcriptome analyses to explore how *P. brassicae* affects *Arabidopsis thaliana* in the night during intermediate (14 days after inoculation; DAI) and late (21 DAI) infection. Day-night differences in gene expression were more pronounced in younger rather than older plants in our differential gene expression (DGE) analysis. Consequently, intermediate phases of infection showed more day-night differences than later ones. Clustering of differentially expressed genes (DEGs) in functional categories highlighted how some of the typical processes known to be disrupted by clubroot infection are more significantly affected in the night and also uncovered some disrupted exclusively in the night. RNA modification stood out as the most unambiguously upregulated process in infected *Arabidopsis* roots in the night. Analysis of the interaction between clubroot infection and diel oscillations in gene expression detected modifications in the rhythmicity of central circadian clock components during the infection. We discuss our findings in the context of manipulation of plant defence and metabolism, identifying targets for experimental validation and highlighting potential new lines of investigation of our time-resolved datasets to better understand the interaction between *P. brassicae* and its host.

**Significance statement:** *Plasmodiophora brassicae* impacts physiological processes under strong temporal control in Brassicaceae hosts: e.g., metabolism, hormone homeostasis and defence. Here, for the first time, we performed a time-resolved transcriptomic exploration of clubroot disease at intermediate and mature stages of infection. We identify a previously unrecognised role for nocturnal manipulation of organellar RNA editing and disruption of rhythmicity in circadian clock components. We provide a dataset enabling further exploration of the impact of clubroot on plant circadian processes.

## Introduction

Responsible for around 10 % of harvest losses worldwide, *Plasmodiophora brassicae* Woronin is one of the most impactful pathogens of Brassicaceae crops (Dixon, 2009). This obligate biotrophic parasitic protist belonging to the class Phytomyxea (Rhizaria; Sierra et al., 2016) has a complex life cycle featuring a short-lasting, symptomless primary phase occurring in the root hairs and epidermis of a broad array of plant hosts; and a secondary, long-lasting and symptomatic phase, specifically occurring in hosts within the family Brassicaceae where durable resting spores are formed (Kageyama & Asano, 2009; Liu et al., 2020). The secondary phase of the infection occurs in the root cortex and stele, where it impacts the host physiology and anatomy, causing successive hyperplasia and hypertrophy in the affected tissues (Olszak et al., 2019), leading to the typical gall formation from which the common name “clubroot disease” derives. This modification of the host anatomy and physiology is required for the parasite to generate a strong local nutrient sink at the site of infection, inducing a systemic redirection of the host photosynthates (Malinowski et al., 2019). Clubroot impact on susceptible plants is heavy, with infected hosts featuring stunted growth, increased susceptibility to wilting and early onset of senescence (Javed et al., 2023). Molecular investigations have been proven paramount in understanding the influence of *P. brassicae* on its host physiology (Agarwal et al., 2011; Ciaghi et al., 2019; Irani et al., 2018; Siemens et al., 2006), highlighting highly impacted processes deserving more extensive investigation, such as: cell wall expansion, remodelling and lignification (Stefanowicz et al., 2021; Tu et al., 2024); cell cycle activation and modification (Olszak et al., 2019) and sugar homeostasis (Siemens et al., 2011). Pathogen-induced disruption of sugar homeostasis is crucial to the trophic interaction between *P. brassicae* and its host, whereby the parasite generates a sugar sink in the gall tissue by increasing symplastic sugar delivery via extracellular invertases (Siemens et al., 2011) and by promoting phloem differentiation and local accumulation of sugar transporters at the site of infection (Walerowski et al., 2018).

Because of its central role in plant physiology, the disruption of sugar metabolism likely mediates systemic reactions at the organism level in clubroot disease, but the substantial anatomical changes observed are also the result of its impact on hormone homeostasis (Devos et al., 2006; Ludwig-Müller et al., 2009). Hormones involved in plant defence such as salicylic acid (SA) and jasmonic acid (JA) have been broadly linked to heightened resistance against *P. brassicae* (Gravot et al., 2012; Lemarié et al., 2015; Lovelock et al., 2013, 2016); and the parasite seems to directly disrupt both (Ludwig-Müller et al., 2015; Smolko et al., 2024). *P. brassicae* also induces accumulation of the stress-related hormone abscisic acid (ABA) as a consequence of the impaired vascular transport and water supply caused by gall formation (Ludwig-Müller et al., 2009). *Arabidopsis* mutants impaired in the perception of the stress-related hormone ethylene (ET) showed increased resistance only when the mutation targeted the crosstalk between ethylene and auxin transport (Ludwig-Müller et al., 2009; Vandenbussche et al., 2003). Indeed, auxins have been found to play an important role in clubroot disease, gradually increasing in the galls with the development of the infection and mutants impaired in indole acetic acid (IAA) biosynthesis were found to produce smaller or developmentally delayed galls (Grsic-Rausch et al., 2000; Neuhaus et al., 2000). Investigations highlighted how the *P. brassicae* protein PbGH3 can conjugate auxins (Smolko et al., 2024) and the recently identified gene *SYNERGISTIC ON AUXIN AND CYTOKININ 1* (*SYAC1*), a cross-talk component between the auxin and cytokinin pathways in *A. thaliana* roots, made plants more susceptible to infection and exacerbated the symptoms of clubroot when constitutively expressed; supporting a pivotal role of the auxin-cytokinin balance in this interaction (Hurný et al., 2020). In biotrophic interactions between plants and pathogens, the role of cytokinins in establishing local nutrient sinks and in suppressing cell death has long been recognized (Walters & McRoberts, 2006), nonetheless the overall impact of *P. brassicae* on host cytokinins is ambiguous. Studies have reported both suppression and increase in metabolism and levels of cytokinins (Dekhuijzen, 1981; Dekhuijzen & Overeem, 1971; Malinowski et al., 2016), but decreased cytokinin degradation on the host side (Siemens et al., 2006) coupled with evidence of endogenous cytokinin production by the pathogen (Müller & Hilgenberg, 1986; Rolfe et al., 2016) seem to support the need for cytokinins in the local establishment of a *de novo* meristematic sink area in the initial phases of infection (Devos et al., 2005, 2006).

The above listed processes impacted by *P. brassicae* are known to be strongly influenced by diel (i.e., day and night) and circadian oscillations in plants. While most living organisms have evolved a way to synchronise their physiology to the environmental rhythms in order to maximize their fitness, plants are particularly affected by these rhythms as they rely for growth and development on light, arguably one of the most rhythmic timers/*zeitgebers* on the planet (Henriques et al., 2018; Sanchez & Kay, 2016). Changes in light and other oscillating environmental parameters such as temperature, entrain the core circadian clock, a transcriptional/translational feedback loop central to the adjustment of rhythmic physiological responses in plants (S. Wang et al., 2022). Moreover, the same rhythmic physiological processes influenced by environmental factors via the circadian clock also act back on it as endogenous *zeitgebers* such as in the case of sugars derived from photosynthetic carbon fixation, whose correlation with the availability of diurnal light has long been ascertained (Graf et al., 2010; Mora-García et al., 2017).

Besides being under strong temporal control, the availability of photosynthates is also spatially resolved. For example, a sucrose transporter involved in phloem unloading has been found to be specifically upregulated in roots during the night, matching a steady increase in nocturnal root growth that peaks shortly after dawn (Durand et al., 2018; Yazdanbakhsh et al., 2011; Yazdanbakhsh & Fisahn, 2010). The synthesis and accumulation of defence hormones has also been reported to be under the influence of diel oscillations in gene expression, in particular the synthesis and accumulation of competing JA and SA; with JA-mediated resistance to herbivores and necrotrophic pathogens being upregulated in the day, whilst SA-mediated resistance to biotrophic parasites heightened in the night (Goodspeed et al., 2012). As already mentioned, the majority of the processes under circadian control also provide retroactive feedback to inform and entrain the circadian clock itself (S. Wang et al., 2022). Since pathogens are known to manipulate hormones and to strongly affect energy homeostasis, it is possible that these processes are central to mediate pathogen-induced modifications on the host internal clock (Lu et al., 2017).

Many biotrophic parasites have been shown to possess intrinsic transcriptionally-mediated rhythms (Rijo-Ferreira et al., 2017, 2020) and in well-investigated pathosystems, such as *Plasmodium* spp. in vertebrates, the parasite contemporarily synchronizes with (Motta et al., 2023) and disrupts (Prior et al., 2019) the circadian rhythmicity of the host.

We reasoned that, as an obligate biotrophic intracellular parasite, *P. brassicae* is likely to have evolved similar ways to influence the rhythmic gene expression of its host in order to maximize its benefits and hinder defence; especially considering its strong influence on host diel processes such as for example carbohydrates allocation to the root (Graf et al., 2010; Malinowski et al., 2019; Yazdanbakhsh et al., 2011). Nevertheless, until now only the diurnal phase of clubroot infection has been explored. This study aims at providing the first time-resolved RNA-seq dataset, covering both day and night, for intermediate (14 days post-inoculation) and mature (21 days post-inoculation) clubroot disease in *Arabidopsis thaliana.* We provide a first exploration of the data generated in order to highlight processes of the host that are influenced by the parasite specifically during the night and further to understand the interplay between diurnal/nocturnal changes in gene expression and clubroot disease.

## Results

### Overview of gene expression levels in infected and healthy plants in the day and in the night

The major factor influencing the clustering of DEGs of *A. thaliana* in PCAs is their response to *P. brassicae* (Fig. 1 A & C). In the intermediate phases of infection (14 DAI) the first two principal components (PCs) explain a cumulative 78% of the variation (Fig. 1 A). Infected and healthy plants strongly separate along the PC1, explaining more than half of the variation in gene expression (63%). Noticeably, the PC2 (explaining 15% of the variation) mostly separated the samples according to night or day (factor “Time point”). Samples collected at 21 DAI seem to be less affected by the time of sampling, with PC1 explaining 83% of the variation and being clearly linked to the infected or healthy status of the plant samples, and PC2 explaining only 5% of the variation (Fig. 1 C) without clear separation of the samples along the factor “Time point”. A PCA of the gene expression levels in uninfected plants resulted in a similar pattern, whereby older plants did not show a time-dependent response (Additional File 3). Befittingly, in plants 14 DAI more than one third (36,75%) of the differentially expressed genes (DEGs) are specific to the night phase and 16,40% to the day (Fig. 1 B), whilst the proportion of DEGs that are exclusively detected during day or night is decreasing at 21 DAI indicating a reduced influence of the factor “Time point” on gene expression from the intermediate to the late stages of infection (21,48% in the night, 17,52% in the day and 61% shared; Fig. 1 D).

**Fig. 1.**
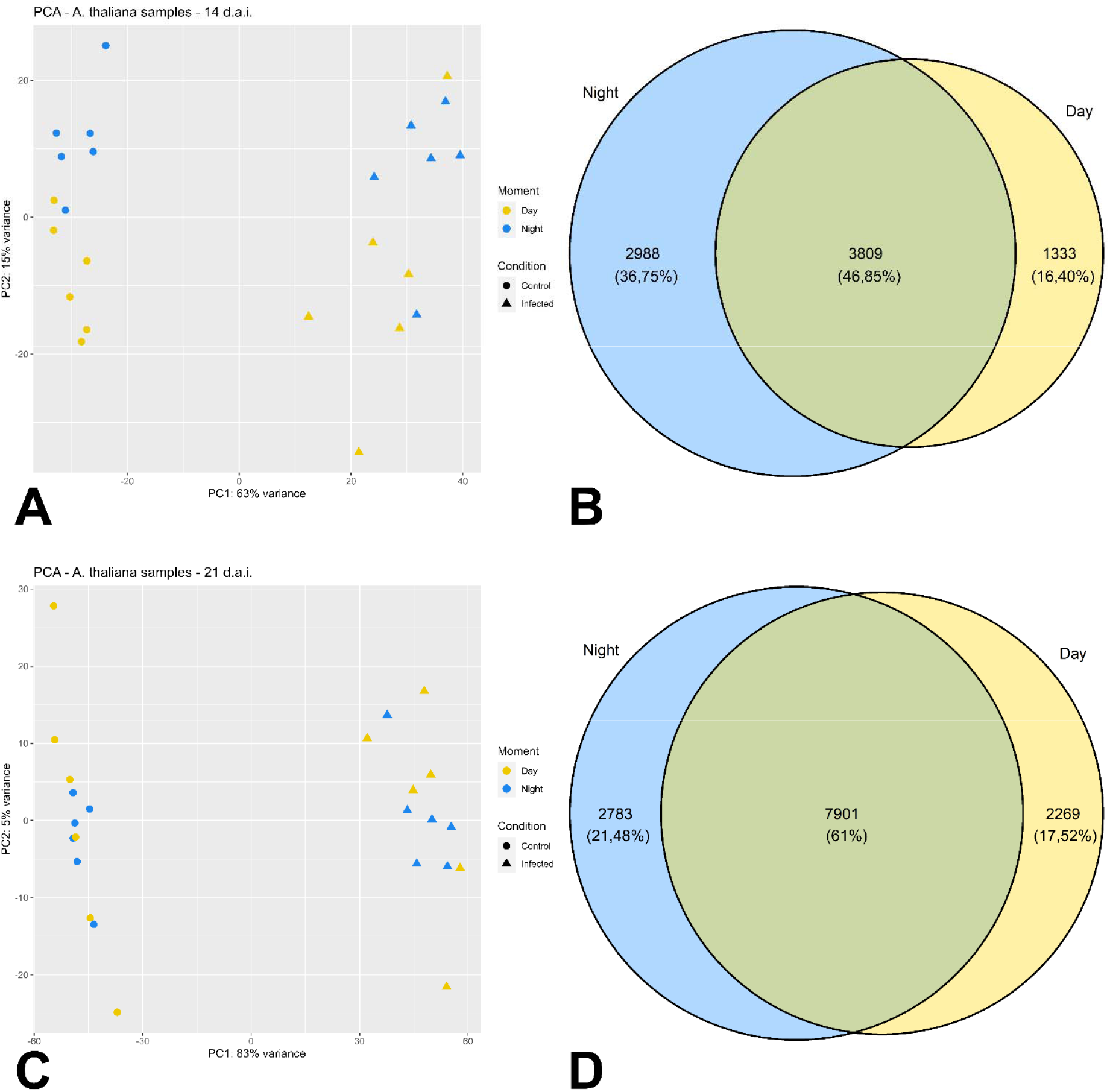
Day/night-informed overview of gene expression in samples of A. thaliana roots 14 DAI (A, B) and 21 DAI (C, D) with P. brassicae. A and C, PCA of A. thaliana root samples structured by gene expression estimates (i.e., vst counts). B and D, Proportions of DE genes between infected and healthy A. thaliana roots assigned to the day, night or both.

### Regulation of enriched GO categories in infected plants during day and night

Differentially expressed genes (DEGs) were largely downregulated in infected plants across all sampling times and DAI, as shown in Additional Files 4 and 5. This overall trend was further confirmed through an analysis of average expression levels within overrepresented Gene Ontology (GO) categories, which revealed a consistent predominance of downregulated over upregulated categories.

At 14 DAI during the day, this imbalance was particularly evident: 63 GO categories showed downregulation compared to just 25 that were upregulated, while one category— “response to molecule of fungal origin”—exhibited an equal mix of both (Fig. 2A and Additional File 6).

**Fig. 2.**
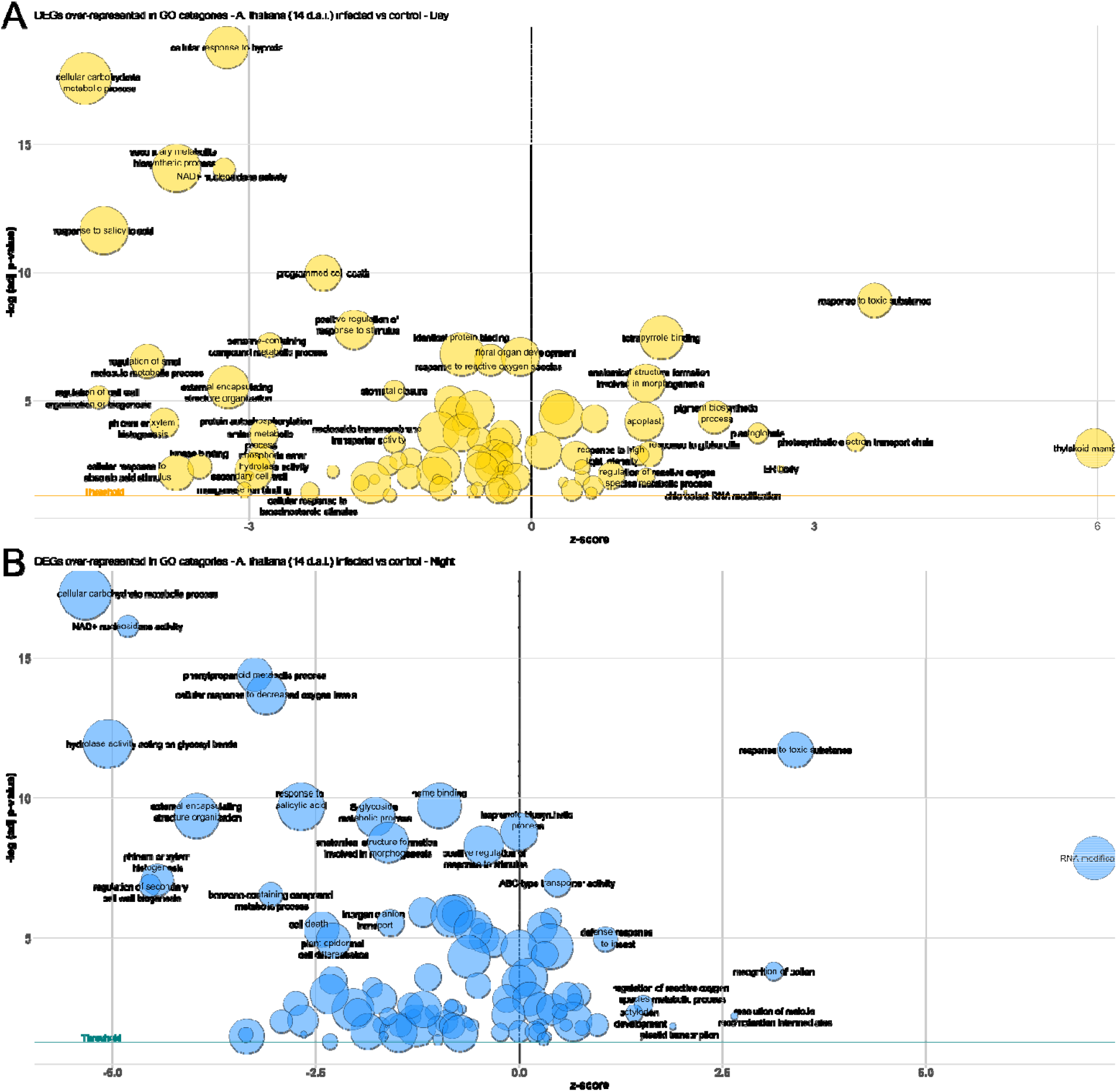
GO bubble chart of GO categories enriched in DEGs in infected A. thaliana roots during the day (A, yellow) and night (B, blue) 14 DAI by P. brassicae. The bubbles are arranged according to their overall expression level or z-score on the x axis and according to the negative logarithmic scale of the adjusted p-value on the y axis. The bubble size is proportional to the gene counts within each category. The significance threshold of adjusted p-value < 0,05 (1,3 in -log10 scale) is represented by coloured lines parallel to the x axis (orange in the day and turquoise in the night).

The ten most significantly upregulated categories during the day were associated with responses to environmental stress, development, and biosynthesis, with highly significant enrichment seen in processes such as response to toxic substances, anatomical structure morphogenesis, pigment biosynthesis, and photosynthetic activity. In contrast, the most significantly downregulated categories were related to core metabolic and defense responses, including hypoxia response, carbohydrate metabolism, salicylic acid signaling, and floral development (Fig. 2 A and Additional File 6).

A similar expression pattern was observed during the night at 14 DAI, with 67 GO categories downregulated and 30 upregulated, alongside five showing no overall change (Fig. 2B and Additional File 6). Upregulated categories at night included many processes related to stress-response and signaling such as response to toxic substance, RNA modification, ABC-type transporter activity, and response to reactive oxygen species. Meanwhile, downregulated categories again pointed to a suppression of key metabolic and defense functions, including phenylpropanoid metabolism, oxygen response, glycoside metabolism, and response to salicylic acid.

By 21 DAI, the contrast between down- and upregulated categories became even more pronounced, consistent with the higher number of DEGs observed at this later stage of infection (Fig. 3, Additional Files 5 and 9). During the day, 94 categories were downregulated compared to just 15 upregulated. These upregulated categories continued to reflect photosynthetic and light-responsive processes, such as plastid organization, pigment biosynthesis, circadian rhythm regulation, and peroxisomal activity. Conversely, the most downregulated categories included pathways related to toxic substance response, carbohydrate metabolism, extracellular signaling, and salicylic acid response, indicating a persistent suppression of metabolic and defense processes.

**Fig. 3.**
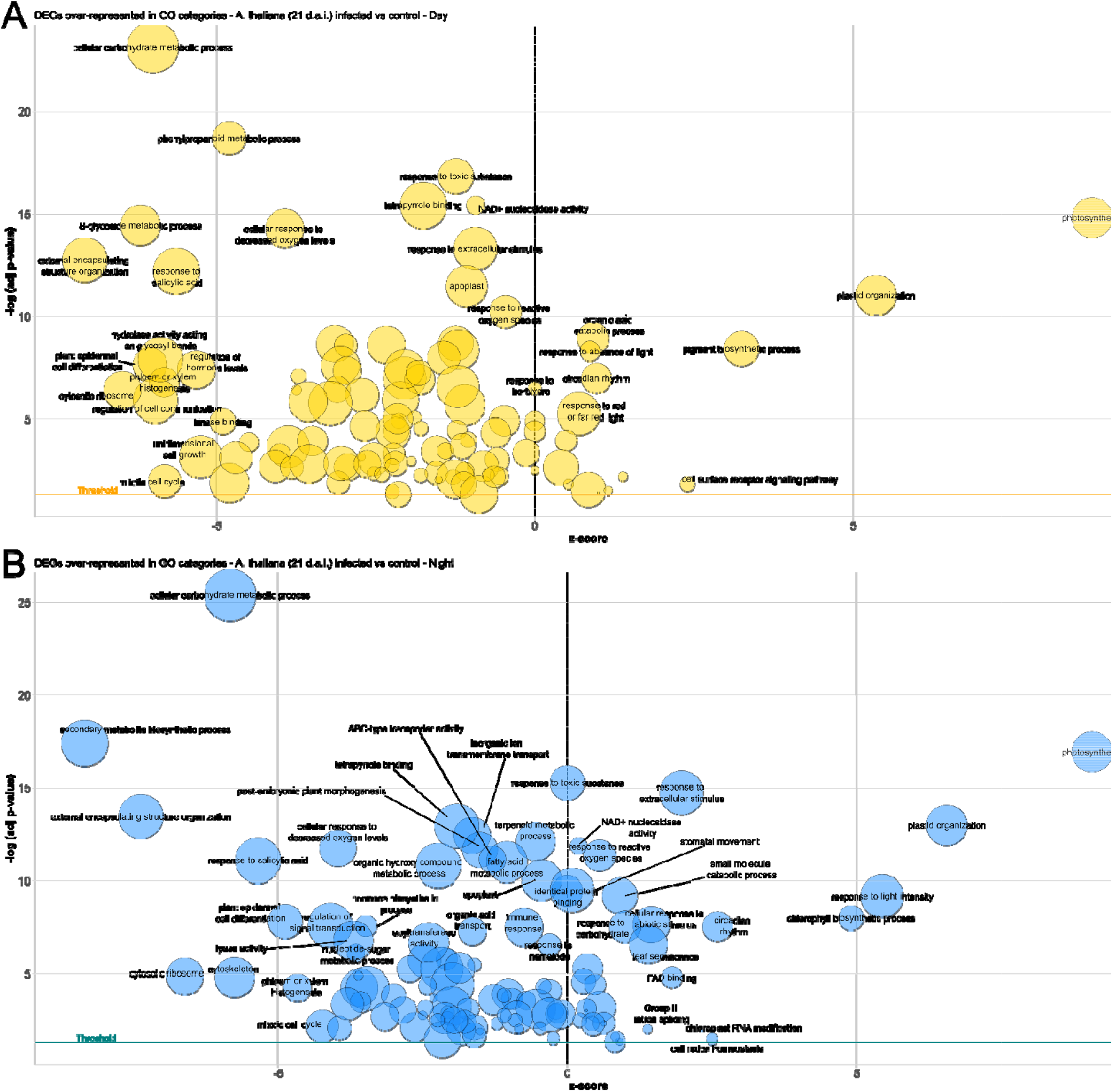
GO bubble chart of GO categories enriched in DEGs in infected A. thaliana roots during the day (top, yellow) and night (blue, bottom) 21 DAI by P. brassicae. The bubbles are arranged according to their overall expression level or z-score on the x axis and according to the negative logarithmic scale of the adjusted p-value on the y axis. The bubble size is proportional to the gene counts within each category. The significance threshold of adjusted p-value < 0,05 (1,3 in -log10 scale) is represented by coloured lines parallel to the x axis (orange in the day and turquoise in the night).

At night, the upregulated GO categories again included several also enriched during the day, such as photosynthesis and plastid organization. Additional night-specific upregulated processes included responses to light intensity, chlorophyll biosynthesis, and abiotic stress. Downregulated categories at night often mirrored those observed during the day, particularly in areas such as carbohydrate metabolism, tetrapyrrole binding, and external encapsulating structure organization. However, some processes—such as inorganic ion transport and secondary metabolite biosynthesis—were uniquely or more strongly suppressed during the night phase.

All GO terms and their associated gene lists used prior to plotting in Figures 2 and 3 are provided in tabular format in Additional File 10.

### Effect of the interaction between clubroot and time of the day on A. thaliana gene expression

The variables “Condition” and “Time Point” are not fully independent, both influence gene expression at the same time. To identify the genes that are affected by both of these variables we used a statistical “interaction effect” model (IE; Duda et al., 2023). This identified 217 functionally annotated genes at 14 DAI which were strongly impacted by both variables (Fig. 4). These 217 DEGs can be grouped in eight clusters according to their expression patterns. Enriched GO terms could be detected in 6 of these clusters (cluster 1, 3, 4, 5, 6 and 8) hinting at how infection and time of the day are affecting processes in the host (Additional File 8 GO terms overrepresentation analysis of clusters of DEGs driven by the interaction between infection and time of the day 14 DAI8).

**Fig. 4.**
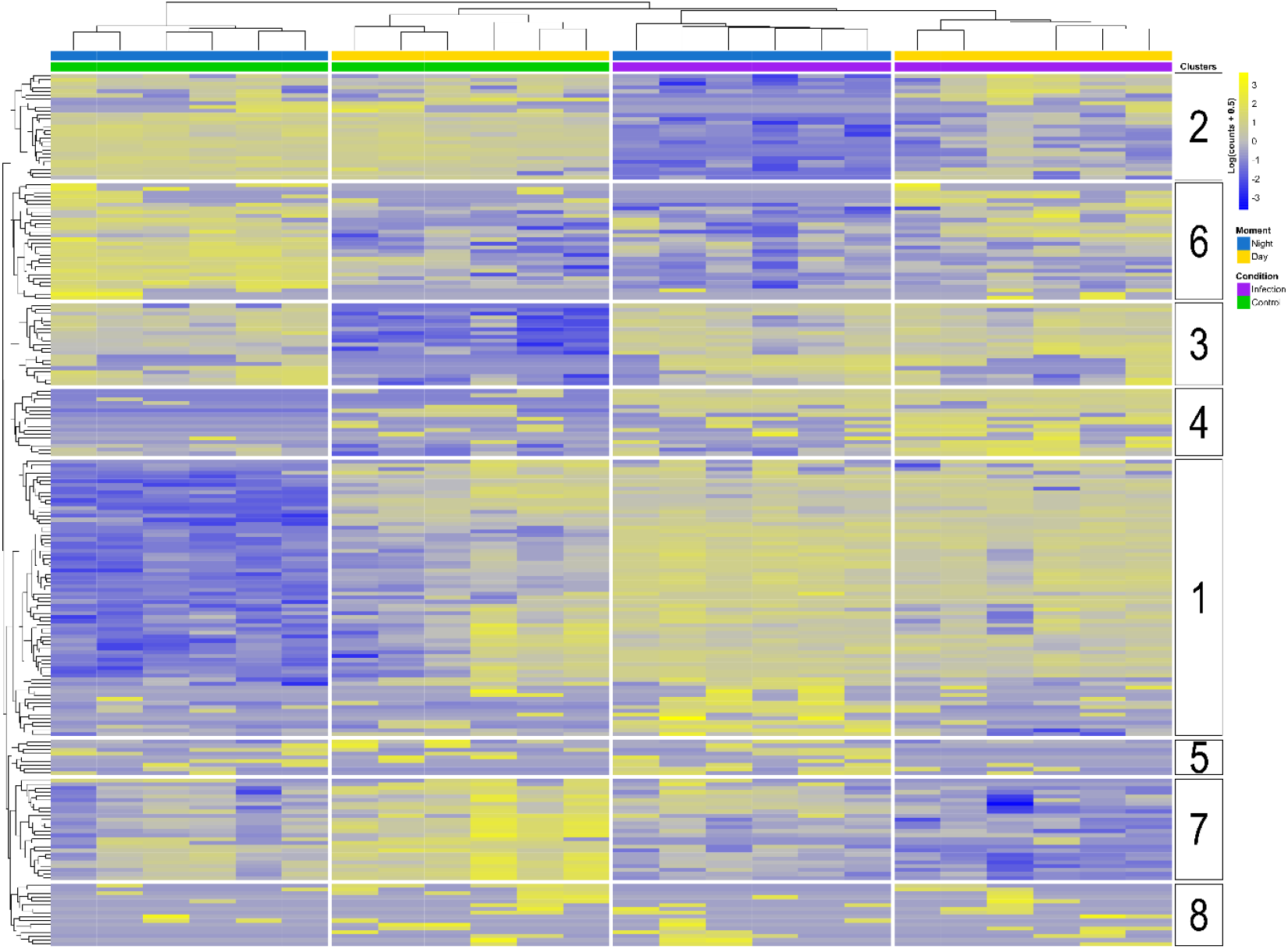
Heatmap highlighting DEGs explained by the interaction between infection and time of the day as detected in infected A.thaliana roots 14 DAI The row-normalised and log transformed gene counts are shown for each sample separately. The status of each sample is described by a combination of green (control plant) or purple (infected plant) and yellow (day) or (blue) bars. DEGs with similar patterns of expression have been grouped in 8 clusters (numbers in white boxes).

Clusters 1 and 3 are characterised by a loss of diel oscillation in infected plants, where the genes are always active regardless of the time of the day. Cluster 1 is enriched in the terms related to senescence, circadian rhythm and cell signalling, where control plants show upregulation during the day. Cluster 3 is characterised by the terms “response to nitrate” (particularly enriched in genes encoding glutaredoxins or GRXS; Ehrary et al., 2020) and terms related to the chloroplast membrane (enriched in genes encoding chlorophyllide a oxygenase, chloro-ribosomal subunits and FKBP16-2), terms showing upregulation in the night in the control. Cluster 7 also loses diel oscillation, but the changes in this cluster are characterised by a downregulation of genes in infected plants which are upregulated during day in control plants.

In contrast cluster 2 acquires a diel oscillation in infected plants with genes being upregulated in the day and downregulated in the night, an oscillation absent in the control plants. Genes in cluster 6 show an inversion of the normal pattern of expression, whereby nocturnal upregulation in the control is overridden in infected plants by a nocturnal downregulation (and vice versa). Cluster 6 is enriched in term “response to fructose”. Cluster 4, enriched in terms related to glycosylation of proteins, shows upregulation by the infection with a stronger effect during the day. Clusters 5 and 8 show no distinct pattern. Cluster 5 is associated with the terms “phosphatidylinositol phosphate kinase activity”, “phosphotransferase activity, phosphate group as acceptor” and “nucleoside transmembrane transporter activity” and cluster 8 with terms related to cellular response to abiotic stress and protein localization to the nucleus.

Fewer genes were impacted by the interaction between time of day and infection at 21 days post-infection (DAI) compared to 14 DAI (96 genes with functional annotations). The clustering of the samples cannot discriminate day and night but it maintains a clear separation along infected and control plants (Fig. 5). The pattern of gene expression could divide the DEGs in five clusters, three of which contain enriched GO terms (Additional File 9). Genes in cluster 1 and cluster 2 show a general upregulation in infected plants and in cluster 2 terms related to response to hypoxia and sugars were enriched. Cluster 3 shows an opposite pattern, with genes mainly downregulated by the infection enriching categories related to the development of anatomical structures and cytoplasmic stress granules. The main expression pattern identified in cluster 4 is the upregulation of genes normally downregulated during the night. This cluster is described by GO terms related to circadian/rhythmic processes, chloroplast membranes, histone methylation and transport of organic acids. Cluster 5 shows a more complex expression pattern, were most of the genes upregulated in the night in control conditions are upregulated in the night in infected conditions and vice versa, but no enriched GO terms were found for this cluster.

**Fig. 5.**
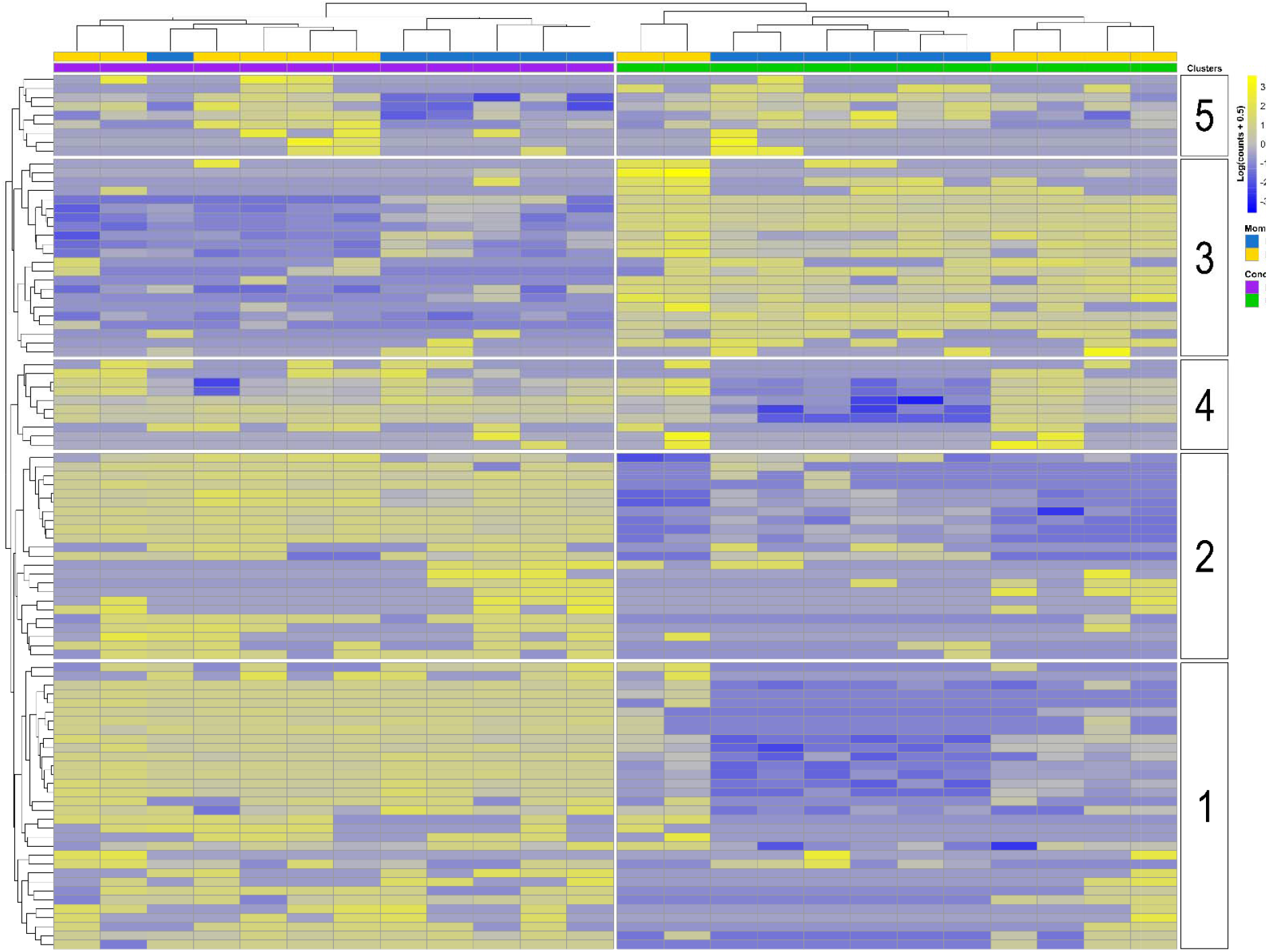
Heatmap highlighting DEGs explained by the interaction between infection and time of the day as detected in infected A.thaliana roots 21 DAI The row-normalised and log transformed gene counts are shown for each sample separately. The status of each sample is described by a combination of green (control plant) or purple (infected plant) and yellow (day) or (blue) bars. DEGs with similar patterns of expression have been grouped in 5 clusters (numbers in white boxes).

## Discussion

This study explores the effects of *P. brassicae* infection on transcriptional diel oscillations in *A. thaliana*, identifying significant effects on gene expression dynamics. Intermediate infection stages (14 DAI) show strong diel influence, aligning with peak metabolic activity of the parasite and initiation of gall formation (Devos et al., 2005; Liu et al., 2020), whereas later stages (21 DAI) reveal reduced rhythmicity, likely driven by plant senescence rather than the pathogen. Nonetheless, overall the strongest driver of gene expression changes is the infection itself, overriding time-of-day effects, and highlighting a complex interaction between host circadian mechanisms and pathogen strategies. Processes such as phenylpropanoid metabolism and RNA modification are disproportionately affected at night and we link these to disruptions of organellar dynamics and defense in infected roots.

Time-resolved RNA-seq analysis of clubroot disease in *A. thaliana* shows an increasing number of DEGs as the infection advances, reflecting the progressive cell-by-cell colonisation of roots by *P. brassicae* (Kunkel, 1918) and its systemic and diversified impact on the plant, where each cell in infected tissues may contain parasitic plasmodia at different stages of development. Existing transcriptomic datasets featuring a developmental dimension support this (Irani et al., 2018; Schuller et al., 2014). The effect of diel changes in gene expression in infected plants appear to be stronger at 14 DAI as compared to 21 DAI (Fig.1) and a decreased influence of day and night on gene expression is also evident in control plants (Additional File 3) indicating a general circadian dampening with plant age. The relationship between senescence and circadian rhythm in ageing plants has been studied in leaves, revealing a similar pattern of reduced amplitude and an increase in frequency of circadian oscillations with age (Kim & Hong, 2019; Lee et al., 2021; Song et al., 2018). A specific circadian clock or ‘root clock’ has been shown to integrate exogenous cues such as light, water availability and temperature in order to adjust root gene oscillations with the external environment (Bustillo-Avendaño et al., 2022). The root clock is closely linked to shoot apical meristem circadian oscillations (Davis et al., 2022; Mortada et al., 2024; Perez-Garcia et al., 2022) and likely influenced by senescence in above ground tissues, although specific work on the root developmental clock has not yet addressed its interplay with root senescence itself (Siqueira et al., 2022). At 21 DAI clubroot infected *A. thaliana* plants are four weeks old and therefore approaching the end of their six-week life-cycle (Meinke et al., 1998). This advanced senescence would explain the reduced influence of diel rhythmicity observed at 21 DAI (Fig. 1 C&D, 3 and 5). In contrast, a clear influence of night and day has been observed at 14 DAI (14 DAI; Fig. 1 A&B, 2 and 4).

The infection is the strongest driver of changes in gene expression (Fig. 1 A), but the genes affected differ between day and night especially in the intermediate phases (i.e., 14 DAI; Fig. 2 and 3; Additional Files 6 and 7). GO categories enriched by infection-related DEGs during the day are in line with previous, comparable studies (Ciaghi et al., 2019; Irani et al., 2018). Nonetheless, interpreting these results requires careful consideration, since each enriched GO category contains upregulated and downregulated genes and since the z-score reflects only the count of genes up/downregulated but not the magnitude of changes. For example, specific genes involved in low oxygen responses during clubroot such as PYRUVATE DECARBOXYLASE 1 (PDC1) and ALCOHOL DEHYDROGENASE 1 (ADH1) have been found to be upregulated in previous studies (Gravot et al., 2016). These specific genes are also upregulated in our dataset, even though low oxygen response categories are overall downregulated (Fig. 2). Similarly, cell wall-related categories are consistently downregulated in our dataset, though individual genes like expansins and pectin methylesterases show contrasting gene-specific expression patterns, which is in line with previous findings (Ciaghi et al., 2019; Irani et al., 2018; Stefanowicz et al., 2021).

The GO term “RNA modification” was the most striking night-exclusive category in terms of differential expression, gene counts and significance at 14 DAI. The majority of genes (90/101) are assigned to pentatricopeptide repeat-containing proteins (PPR proteins), with most of them being upregulated (86/101) and part of the PLS-class PPR proteins (67/101). Land plant genomes harbour an expanded number of PPR protein encoding genes, specifically PLS (Barkan & Small, 2014), and many of them are involved in the RNA stabilisation, splicing and editing in chloroplasts and mitochondria, necessary for the correct development and functioning of the target organelle (Barkan & Small, 2014; X. Wang et al., 2021). These data thus suggest a pivotal role for nocturnal manipulation of mitochondria and plastids by *P. brassicae.*

On the one hand, the upregulation of genes potentially involved in organelle dynamics in infected *A. thaliana* would fit the well-documented phenomenon of accumulation of amyloplasts in the root cortex of plants affected by clubroot (Garvetto et al., 2023; Ma et al., 2022). Disruption of transcription is known to inhibit the differentiation of plastids into amyloplasts (Enami et al., 2011; Liebers et al., 2017), suggesting a potential involvement of post-transcriptional editing in the transition. Furthermore, cytokinin concentration, another factor necessary for the transition to amyloplast (Enami et al., 2011), is increased locally by *P. brassicae* (Müller & Hilgenberg, 1986; Siemens et al., 2006) arguing for its importance in clubroot development. Amyloplast induction driven by alterations in the cytokinins-to-auxins ratio would enable the storage of sugars directed toward the rapidly growing root in the night (Graf et al., 2010; Yazdanbakhsh et al., 2011; Yazdanbakhsh & Fisahn, 2010), where the parasite generates an even stronger physiological sink (Keen & Williams, 1969; Malinowski et al., 2019; Walerowski et al., 2018). Nevertheless, on the other hand, plastidial RNA editing has been found to be more effective in green tissues as compared to non-photosynthetic ones (Peeters & Hanson, 2002) and a high proportion of PPR in our data seem to target the mitochondrion. Therefore, a hypothesis encompassing organelle-mediated defences might be more parsimonious.

RNA editing is required for the correct functioning of chloroplasts and mitochondria and mutations of PPR proteins often lead to defective growth or lethal phenotypes, in particular by stimulating the production of reactive oxygen species (ROS) due to defects in the post-transcriptional modifications of genes targeting these two organelles (Y. Wang & Tan, 2025). Evidence exists of plants exploiting increased ROS levels caused by the disruption of the electron transport chain in chloroplasts and mitochondria as a signal to promote defence or to initiate oxidative burst and fend off pathogens. In *A. thaliana* chloroplasts, reduced editing of *ndhB* mRNA (encoding the B subunit of the chloroplast NADH dehydrogenase-like complex -NDH) caused by the disruption of PPR proteins CRR21 and CRR2 (both upregulated in our dataset) led to substantially enhanced ROS production and subsequent resistance to fungal pathogens (García-Andrade et al., 2013). A recent study found that lack of editing and splicing of mitochondria-targeting mRNAs *nad1, nad3, nad6* and *nad7* (encoding subunits of mitochondrial Complex I) caused by downregulation of the PPR protein RESISTANCE TO PHYTOPHTHORA PARASITICA 7 (RTP7) results in ROS overproduction, oxidative burst and incompatible interaction with the pathogenic oomycete *Phytophthora parasitica*(Yang et al., 2022) Although not a PPR, multiple organellar RNA-editing factor 8 (MORF8) participates in RNA editing in both chloroplasts and mitochondria, and has been shown to decrease resistance to *P. parasitica* when targeting the mitochondrion, putatively by stabilizing the respiratory chain thereby avoiding ROS production, SA signalling and oxidative burst (Yang et al., 2020). This is particularly interesting as *morf8* is among the genes assigned to “RNA modification” upregulated in the night in our analysis. Therefore, the increased RNA editing observed in our dataset can be interpreted as an attempt from the side of *P. brassicae* to maintain functional mitochondria and plastids, subsequently dampening ROS production and defence. The hormone salicylic acid (SA) stimulates mitochondrial ROS production (J. Wang et al., 2022) and is involved in a positive feedback loop (Mencia et al., 2020), suppressing mitochondrial RNA editing and being induced by the ensuing ROS production as observed in (Yang et al., 2020). SA is also under strong diel control and peaks in the middle of the night (Zheng et al., 2015; Lu et al., 2017; Karapetyan & Dong, 2018), therefore disruptions in SA signalling by *P. brassicae* in the moment of its normal peak concentration might help to explain the enhanced nocturnal upregulation of “RNA modification” in our analyses.

Indeed, SA-mediated defences are known to be affected in the interaction between *P. brassicae* and its host (Galindo-González et al., 2020; Mencia et al., 2022), as exemplified by the parasite’s effector protein PbBSMT methylating SA to methyl-salicylate (MeSA) in order to enhance its transport away from the site of infection, thereby reducing its local concentration and dampening the immune response (Djavaheri et al., 2019; Ludwig-Müller et al., 2015). Accordingly, an overall downregulation of the category “response to salicylic acid” was observed in our analyses in the intermediate and mature infection regardless of the time of the day (Fig. 2 and 3; Additional File 10).

A putative targeting of nocturnal SA signalling by *P. brassicae* might also explain why the biosynthesis of phenylpropanoids, enhanced by SA (J. Y. Chen et al., 2006), is more severely impacted in the night (64 genes enriching the category “phenylpropanoid metabolic process” in the night as compared to 49 in the day; Additional File 10; Bauters et al., 2021). Phenylpropanoids are known to be involved in plant mechanical and chemical defence, especially via the induction of lignin deposition in secondary cell walls and the accumulation of flavonoids (Dong & Lin, 2021). The pivotal importance of lignification of secondary cell walls in preventing the colonization and spread of *P. brassicae* is known, with the parasite actively interfering in this process in susceptible *B. napus* varieties (Tu et al., 2024) and downregulation of lignin biosynthesis has been observed in clubroot infected *A. thaliana* (Agarwal et al., 2011). Our data suggest that the manipulation of the nocturnally gated SA response by *P. brassicae* is a key driver of the changes observed in defence and organellar dynamics, arguing in favour of the exploration of nocturnal phases of clubroot to better understand this disease. Furthermore, the recently established link between SA acid and circadian clock(Fraser et al., 2024) advocates for the identification of genes whose oscillations are equally affected by temporal and disease dynamics.

The IE model allowed for the detection of this small and interesting subset of DEGs whose expression pattern is simultaneously affected by time of the day and infection (Duda et al., 2023). The overall reduction of DEGs at 21 DAI as compared to 14 DAI (i.e., 96 against 217) is likely due to the already observed decreased effect of diel oscillations in older plants and explains the absence of day/night grouping of the samples at 21 DAI (Fig. 5). At 14 DAI, when the effect of time on gene expression is stronger, DEGs appropriately divide samples into day/night and infected/control groups (Fig. 4). At 14 DAI, row clustering reveals varying effects of infection on the diel gene expression patterns, the most notable being the suppression of rhythmicity through upregulation of genes that are normally downregulated at night (i.e., cluster 1 in Fig. 4). Interestingly, the GO term enrichment analysis of the DEGs in cluster 1 features the terms “Circadian rhythm” and “Rhythmic process” (Additional File 8). A closer look at the list of genes contained in cluster 1 revealed seven genes of acknowledged importance in rhythmic and circadian processes: *GIGANTEA* (GI), *PSEUDO-RESPONSE REGULATOR* 3 (PRR3), 7 (PRR7) and 9 (PRR9); *PHYTOCHROME INTERACTING FACTOR 4* (PIF4), *NIGHT LIGHT-INDUCIBLE AND CLOCK-REGULATED* 1 (LNK1) and 4 (LNK4). Members of the core circadian clock such as GI and the PRRs are known to be directly involved in the transcriptional/translational feedback loop central to circadian oscillations (Sanchez & Kay, 2016), but also to integrate endogenous and exogenous stimuli to inform and entrain it (S. Wang et al., 2022), a role they share with PIF4 and LNK1/4 (Kidokoro et al., 2023; Zhu et al., 2016). Whilst LNK1 and LNK4 seems to mainly integrate temperature stimuli in the circadian oscillator (Kidokoro et al., 2023), all other genes detected above are variously involved in the integration of information about temperature, light, sugar and hormone levels within the circadian clock (S. Wang et al., 2022). While temperature and light are unlikely to be involved in mediating the pathogen influence on the diel rhythm of the host, *P. brassicae* influence on hormones (Jayasinghege et al., 2023; Wei et al., 2021) and sugar levels (Keen & Williams, 1969; Ma et al., 2022) is documented and a more plausible mediator. Considering the suppression of the diel cycle observed in cluster 1 and the fact that the majority of the rhythmic genes we identified within it are inhibitors of the central oscillator CIRCADIAN CLOCK ASSOCIATED 1 /LATE ELONGATED HYPOCOTYL (CCA1/LHY), either directly such as PRR9, PRR7 (Farré & Liu, 2013) or indirectly such as PRR3 (via TOC1; Para et al., 2007) and LNK1 (via RVE8 and PRR5; Kidokoro et al., 2023), it is possible that these genes are mediating the influence of the pathogen on the circadian clock.

In the shoot PRR7 is activated in condition of low sugars by a mechanism involving the activation of the sugar sensor SUCROSE NON FERMENTING RELATED KINASE 1 (SnRK1), itself phosphorylating the transcription factor bZIP63 which activates PRR7 transcription (Frank et al., 2018). Interestingly, SnRK1 has been found to be the direct target of the recently described *P. brassicae*-specific effector protein PBZF1 (W. Chen et al., 2021). SnRK1 is a central sensor of nutrient stress in plants (Han et al., 2024), and the fact that this gene seems to be at the center of direct (i.e., via effectors) and indirect (i.e., via changes in sugar levels) manipulation by *P. brassicae* argues for its pivotal role in the interaction. Recent studies have shown that sucrose has an even higher importance in conveying circadian information in the root, where sucrose pulses from the shoot regulate diel oscillations of PRR7 (and PRR9; Uemoto et al., 2023). Unlike sucrose and trehalose-6-phosphate levels, plants do not regulate starch turnover based on its abundance (Seki et al., 2017). As a result, the parasites direct consumption of sucrose and/or the induction of amyloplastic storage could create a continuous flow of photosynthates towards the infection-induced sink, disrupting the normal rhythmic pattern of sugar delivery to the root. This scenario of a nutrient deprived tissue with consistent expression of PRR7 is further supported in our study by the high expression of the “starvation gene” SKIP20 (a.k.a. KMD4 or At3g59940; Graf et al., 2010) in infected roots.

Despite showing expression patterns like the one discussed above, GI and PIF4 are less clearly influenced by sugar metabolism. Sucrose pulses in sugar-depleted condition in the dark are known to be able to re-establish the oscillations of the circadian clock by post-transcriptionally stabilizing GI but they do not change its expression levels (Dalchau et al., 2011; Haydon et al., 2017); whilst sucrose influences the expression pattern of various PIFs, but not of PIF4 (Shor et al., 2017). GI is a plant specific circadian protein promoting the transcription of thousands of genes and whose competitive interaction with PIF4 integrates endogenous oscillations with light and temperature in order to optimize growth (Nohales et al., 2019; Park et al., 2020). Among the many processes influenced by GI, senescence is particularly interesting, since it is also found in our GO term enrichment analysis of cluster 1 (“Plant organ senescence”; “Leaf senescence”; Additional File 8) and prior studies have shown delayed senescence when *GI* expression is suppressed (Thiruvengadam et al., 2015).

In our dataset, the established connection between circadian rhythms and senescence processes may be linked to the continuous expression of *GI* during the night, explaining the premature senescence and the yellowing of leaves observed as typical clubroot symptoms. GI nocturnal breakdown also regulates gibberellin mediated growth by allowing DELLA degradation and enabling the expression of downstream growth promoting genes (Nohales & Kay, 2019). The upregulation of *GI* in the night in our dataset might therefore be involved in disrupting the root growth (especially lateral root formation and main root elongation) by making cells insensitive to gibberellin (Shtin et al., 2022). Intriguingly, GI is also involved in stabilization of REPRESSOR OF GA (RGA; Park et al., 2020), itself inhibiting the gibberellin-induced transcription of PIF4 (another promoter of growth genes), which is instead upregulated in the night in infected plants. These results highlight the complexity of the influence of the circadian oscillator on downstream processes acting at the post-transcriptional level.

## Conclusions

Overall, our exploration highlights a previously unrecognized nocturnal impact of *P. brassicae* on organellar dynamics and plant defences, and suggests a disruption of the host circadian processes. We provide target genes amenable to experimental validation, the necessary next step in order to answer questions on the time-specific cross-talk between *P. brassicae* and its host. Nocturnal manipulation of host organellar RNA editing by the parasite is likely to be linked to the diel oscillations in SA signalling and seemingly mediated by a subset of transcripts enriched in PPR encoding genes. The question remains whether PRR genes are a direct target of *P. brassicae* manipulation or whether the influence of the pathogen on their expression pattern is mediated by manipulation of SA signalling. Ongoing analyses of the diel changes in gene expression on the side of the parasite might help identifying time-specific effectors and help clarifying their targets. Furthermore, our investigation already revealed that circadian oscillations in gene expression (i.e., oscillations at a finer scale than the day and night) are altered by *P. brassicae*. In line with recent findings (Fu et al., 2025), the application of specific tools for the analysis of differential rhythmicity to our dataset may further identify effectors allowing *P. brassicae* to hijack the circadian clock components to disrupt the physiology of the host plant.

## Experimental procedures

### Experimental setup & biological material

After germination, seedlings of *Arabidopsis thaliana* ecotype Col-0 obtained from the Seed Stock Centre (stock N1092, Nottingham, UK) were grown on a soil substrate (Einheitserde Type P, pH 5.8; Hermina Samen, Germany) mixed with sand (Sahara Spielsand; Hornbach Germany) in the ratio 4:1. This soil mixture was steam-sterilized prior to use for 120 minutes at 100 °C. All plants were grown in a Sanyo MLR-350H climate chamber at a temperature of 23°C during the day and 18°C during the night. The light/dark cycle was set to 12:12 hours, with a light intensity of 100 μmol. Genome-sequenced *Plasmodiophora brassicae* isolate e3 (Schwelm et al., 2015) obtained from Dr. Siemens (Fähling et al., 2004) was propagated on *B. rapa* var*. pekinensis* (ISP International Seed Processing GmbH, Quedlinburg, Germany) and pure spore suspensions were obtained as described in (Smolko et al., 2024). One week after germination, *A. thaliana* Col-0 plants were individually inoculated with 1 mL of *P. brassicae* isolate e3 spore suspension at a concentration of 10^6^ spores/mL in 50 mM KH PO buffer (pH 5.5, adjusted with K_2_HPO_4_) or with a mock inoculation of an equal amount of the same buffer. Healthy controls and infected plants were harvested 14 and 21 days after inoculation (DAI, Additional File 1 A and B), in each occasion at four different time points: two points during the “day” (5 hr 50’ and 10 hr 40’ after artificial dawn) and two points during the “night” (15 hr 30’ and 19 hr 20’; i.e., 3 hr 30’ and 7 hr 20’ after artificial dusk). The time point 14 DAI was chosen as it marks the establishment of the secondary infection phase with colonization of cortical tissue by metabolically active plasmodia and onset of visible symptoms in the root (Liu et al., 2020). The second time point (i.e., 21 DAI) has been chosen as it matches the shift from actively feeding plasmodia to resting spore production (Liu et al., 2020). Eight to twelve plants were grown per each time point /treatment and harvested by pooling four plant roots in 1 mL ice-cold RNAlater (Ambion, Austin, TX, USA) in 1.5 mL Eppendorf tubes. Samples were then left to rest overnight, before removing and refreshing the RNAlater and storing them at −80°C until RNA extraction.

### RNA extraction & sequencing

Samples in RNAlater were thawed on ice and one or two plant roots (when biomass from a single root specimen was judged insufficient for RNA extraction) per each tube were transferred to ethanol-sterilised and RNAseAway treated tin foil squares with equally treated steel tweezers (Additional File 1 C and D). A root sample as described above constitute one replicate, and triplicate were selected and processed per each time point, for infected and mock inoculated control plants at 14 DAI and 21 DAI, resulting in a total of 48 processed roots (Additional File 2). Roots were carefully wrapped and snap-frozen in liquid nitrogen before transferring them in pre-mixed bead matrix tubes (D1034-MX; Biozym, Hessisch Oldendorf, Germany) amended with 200 μL IB buffer (SPLIT RNA Extraction Kit, Lexogen, Vienna, Austria) to decrease the risk of RNA degradation and immediately homogenized with a FastPrep bead mill (MP Biomedicals, Santa Ana, CA, USA) for 40 s at 60 ms^-1^ three times. After tissue disruption, another 200 μL of ice cold IB buffer was added to the samples. The RNA extraction proceeded following the protocol SPLIT RNA extraction of total RNA (SPLIT RNA Extraction Kit, Lexogen, Vienna, Austria) as suggested by the manufacturer without further modifications. Assessment of the RNA extracts quality and quantity was performed on an Agilent Bioanalyzer 2100 (Agilent Technologies, Palo Alto, CA, USA) and Quantus fluorometer (Promega Corporation, Fitchburg, WI, USA). Strand-specific library preparation was performed on poly-A-selected mRNA using a NEBNext Ultra II Directional RNA Library Prep Kit with NEBNext Poly(A) mRNA Magnetic Isolation Module and NEBNext Multiplex Oligos for Illumina (New England Biolabs, Ipswich, MA, USA). The quality and concentration of cDNA in the resulting libraries was assessed on an Agilent Bioanalyzer 2100 and Quantus fluorometer, before paired-end sequencing (2×150 bp on a NovaSeq S1; Illumina, San Diego, CA, USA) at the VBCF NGS Unit (Vienna, Austria).

### Sequencing data processing

Read quality was checked using FastQC (Andrews, 2010) and quality reports visualized in MultiQC v1.6 (Ewels et al., 2016) before and after read filtering and adapter trimming with Trim Galore v0.6.2 (quality score threshold 30; https://zenodo.org/doi/10.5281/zenodo.5127898). To assign reads to the host plant and the parasite, Bowtie2 v2.3.5.1 (Langmead & Salzberg, 2012) was used to map them against the concatenated genomes of *Arabidopsis thaliana* (TAIR10; Lamesch et al., 2011) and *Plasmodiophora brassicae* (Pldbra_eH_r1) with a “Place-to-go-based” approach as defined in (O’Keeffe & Jones, 2019). Obtained read pools (separately for the host and the parasite) were assembled with Trinity v2.10.0 (in genome guided mode; Grabherr et al., 2011)) and resulting transcripts annotated with Trinotate v3.2.1 (Bryant et al., 2017). Expression estimates were obtained by pre-aligning the reads to the transcripts using STAR v2.7.1a (Dobin et al., 2013) and assigned to assembled transcript with Salmon (Patro et al., 2017). Raw NGS sequences of the dataset obtained in this study are deposited in NCBI Sequence Read Archive (SRA) under the BioProject accession number PRJNA1212907.

### Differential gene expression

DESeq2 Bioconductor package (Love et al., 2014) in R (R Core Team, 2021) was used to estimate differential gene expression. In order to detect differences in the response of plants to infection by *P. brassicae* separately during the day and during the night, differentially expressed genes (DEGs) were calculated on pooled counts. Briefly, read counts from all time points sampled during the day (i.e., 5 hr 50’ and 10 hr 40’ after artificial dawn) and during the night (15 hr 30’ and 19 hr 20’ after artificial dawn) were pooled before further analyses.

This resulted in six replicates for each of the four conditions, as defined by the two factors “Condition” (Infected or Control) and “Time point” (Day or Night; Additional file 2). An experimental design with two factors and interaction was used for estimating the DEGs separately for 14 DAI and 21 DAI, thus resulting in five comparisons: infected vs control in the day, infected vs control in the night, control in the night vs in the day, infected in the night vs in the day and interaction. Genes were considered differentially expressed when the adjusted p-value was ≤ 0.05 (Wald test p-value adjusted for multiple testing with Benjamini–Hochberg correction) and a log_2_-fold change was ≤ −1 or ≥ 1.

### Principal component analysis and Venn diagram

Principal component analysis (PCA) was calculated in the DESeq2 Bioconductor package in R (Love et al., 2014) and further modified in ggplot2 (Wickham, 2010). The PCA was performed on variance stabilizing transformed (vst) count data from the DESeqDataSet object, without taking into account the sample information (i.e., blind=TRUE), as suggested by the DESeq2 manual. Venn diagrams on DEGs were calculated in R with the Package VennDiagram (H. Chen, 2022).

### GO term over representation analyses

ClusterProfiler (Wu et al., 2021) was used within R to perform gene ontology (GO) over-representation tests using lists of DEGs in the comparisons between infected and healthy plants in the day and in the night. Significance of the enriched GO categories was assessed by Fisher’s exact test controlled for False discovery rate (FDR) with Benjamini–Hochberg correction and the q-value cutoff was set to 0.05. Because the RNA-seq strategy adopted was designed to measure all expressed mRNAs (total RNA extraction followed by poly-A tail selection), the background gene list (or “universe”) against which over-representation was performed, was composed of a list of all protein coding genes from *A. thaliana* TAIR10 genome release. The package GOplot (Walter et al., 2015) was used to combine expression data with functional categories. Using the function “circle_dat”, the package was used to summarise and append to each enriched GO category an overall expression value (i.e., a z-score) calculated as follows:

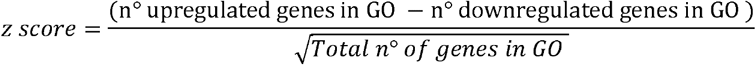

The function “reduce_overlap” within the same package was used to eliminate all but one GO category sharing more than 25% of the genes before plotting the results with the function “GObubble”.

### Heatmap of gene expression levels

Genes whose differential expression levels were driven by the interaction between the factor “Time point” and the factor “Condition” were detected and their expression levels were visualised with heatmaps as implemented in the package pheatmap (Kolde, 2019) in R, on row-scaled, log-normalised read counts and with default “Euclidean” clustering method. All figures were further refined and finalized in Inkscape (Inkscape Project, 2020).

## Author contributions

Experimental concept and design: SN, JLM. Wet lab work: SA, FB, AG, MH. Bioinformatics and statistical analysis: AG. Analysis of Results: AG, SN, SA, JLM, MH. Manuscript writing: AG, SN, SA, JLM. Figures and tables: AG. All authors read and approved the final manuscript.

## Acknowledgements

We are grateful to Bettina Schneidhofer for her support in the wet lab. We would also like to thank the VBCF NGS Unit (www.vbcf.ac.at) in particular Carmen Czepe for her support during the sequencing and Nadine Tatto for her paramount contribution and continuous support with the bioinformatic analyses.

## Data availability statement

Raw NGS sequences of the dataset generated and analysed in the current study are available in NCBI Sequence Read Archive (SRA) under the BioProject accession number PRJNA1212907 or are available from the corresponding author on request.

## Supporting Information

**Additional file 1.** Above and below ground phenotypical traits of infected and control plants at 14 and 21 DAI.

**Additional file 2.** Detailed overview of the samples sequenced in this study.

**Additional file 3.** PCA on control *A. thaliana* plants, showing the decreased influence of the temporal factor with plant age.

**Additional file 4.** Volcano plot showing the number of DEGs in the day and night in infected *A. thaliana* plants 14 DAI.

**Additional file 5.** Volcano plot showing the number of DEGs in the day and night in infected *A. thaliana* plants 21 DAI.

**Additional File 6.** GO categories enriched in DEGs in *A. thaliana* roots 14 DAI infected by *P. brassicae.*

**Additional file 7.** GO categories enriched in DEGs in *A. thaliana* roots 21 DAI infected by *P. brassicae.*

**Additional file 8.** GO terms overrepresentation analysis of clusters of DEGs driven by the interaction between infection and time of the day 14 DAI.

**Additional file 9.** GO terms overrepresentation analysis of clusters of DEGs driven by the interaction between infection and time of the day 21 DAI.

**Additional file 10.** Enriched GO terms and corresponding genes underpinning figure 2 and 3 before data reduction.

## Supporting information

**Additional File 1.**
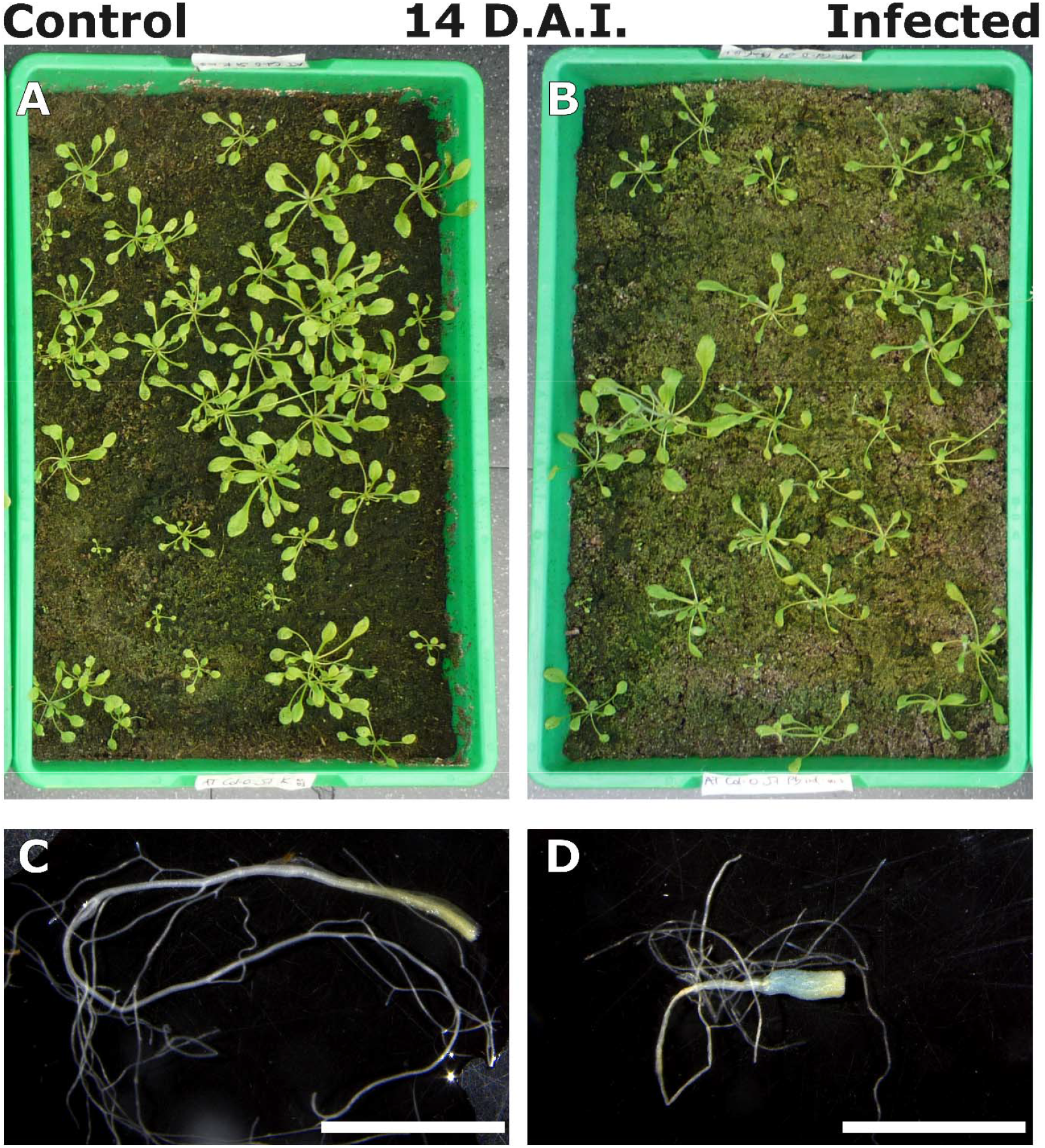
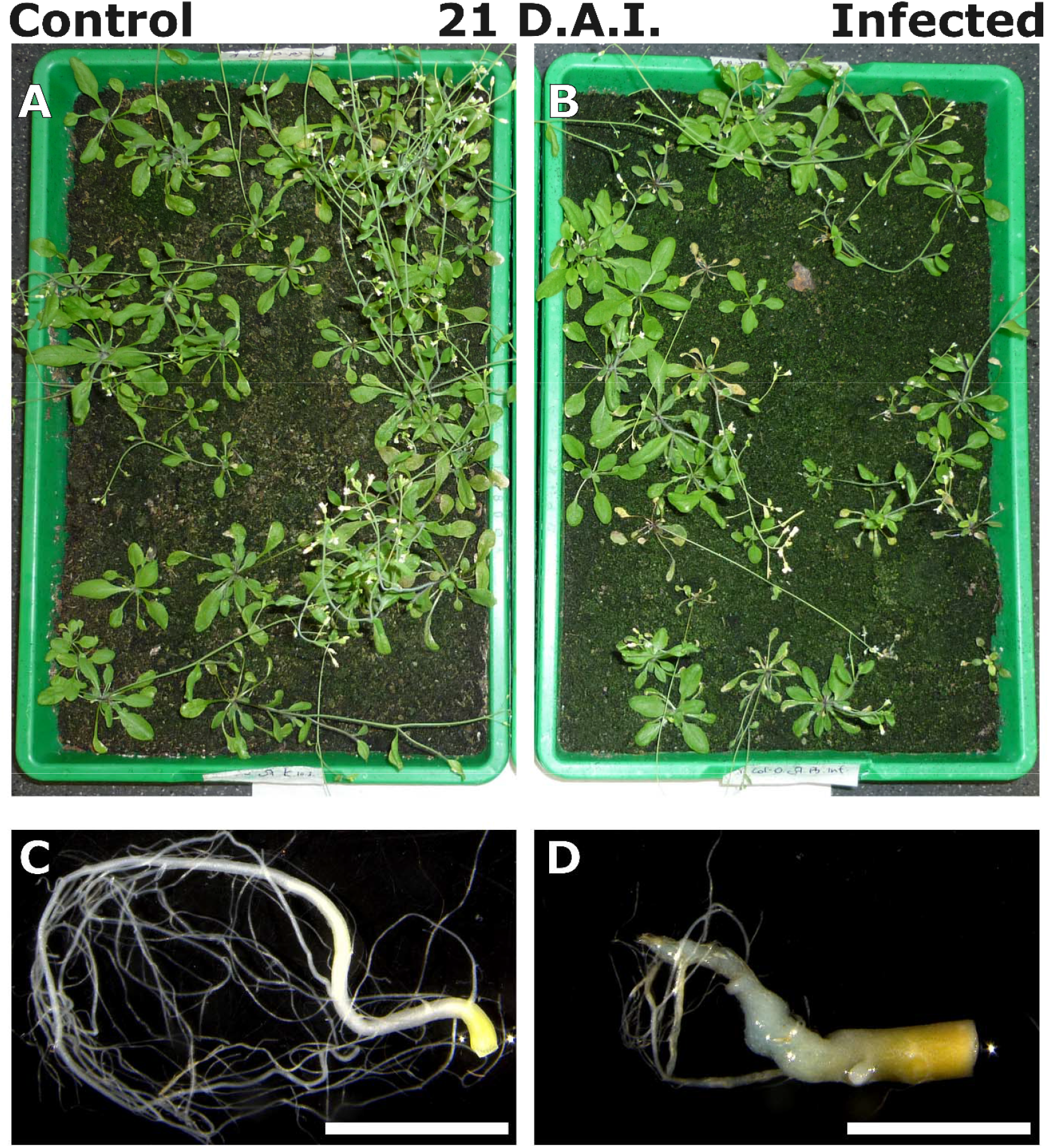
Phenotypical traits of rosettes (A and B) and root samples (C and D) for specimen of healthy and infected A. thaliana Col-0 harvested at 14 d.a.i. (top figure plate) and 21 d.a.i. (bottom figure plate). Scale bars in C and D = 5 mm.

**Additional File 2.**
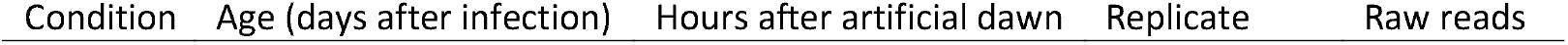
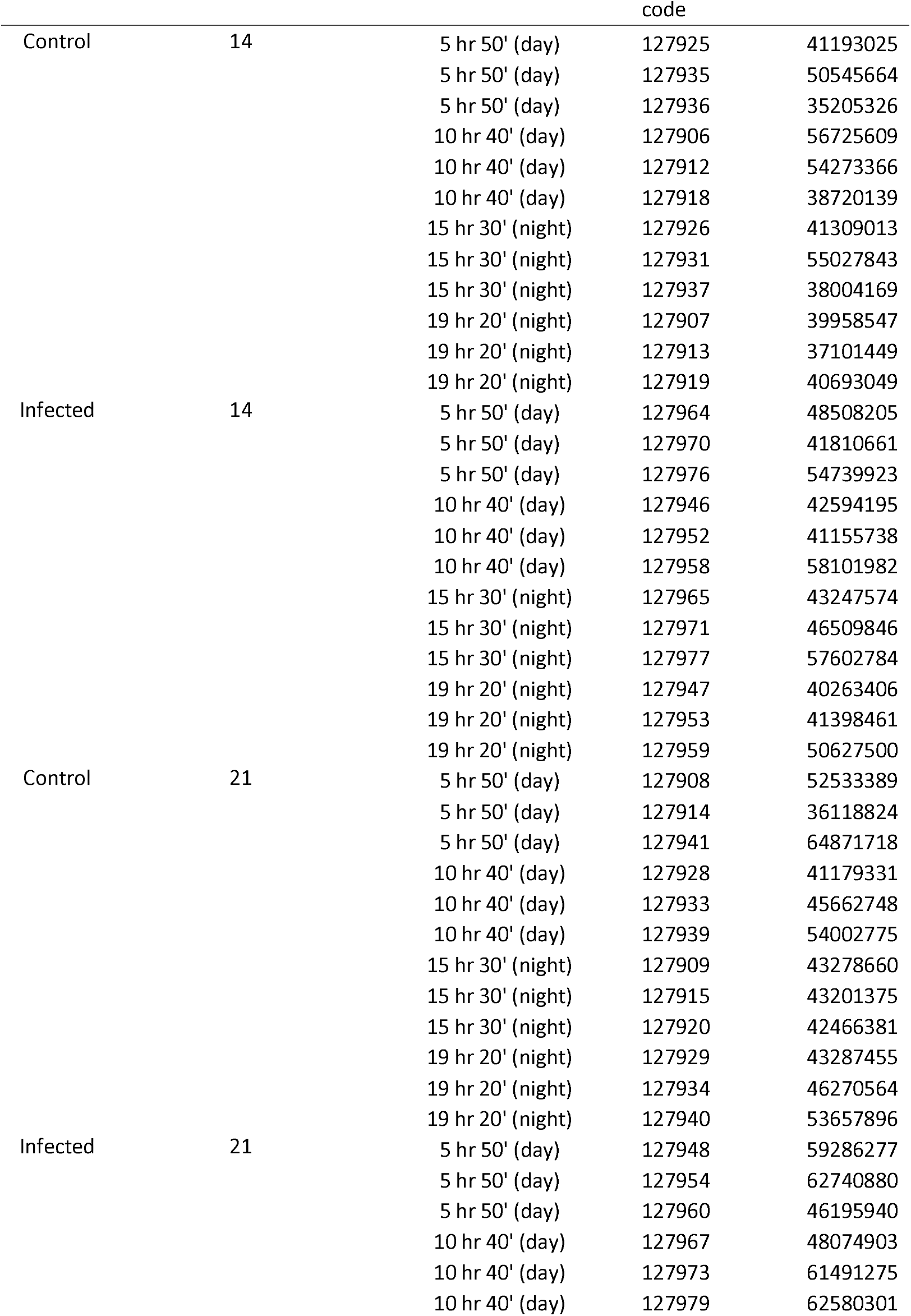
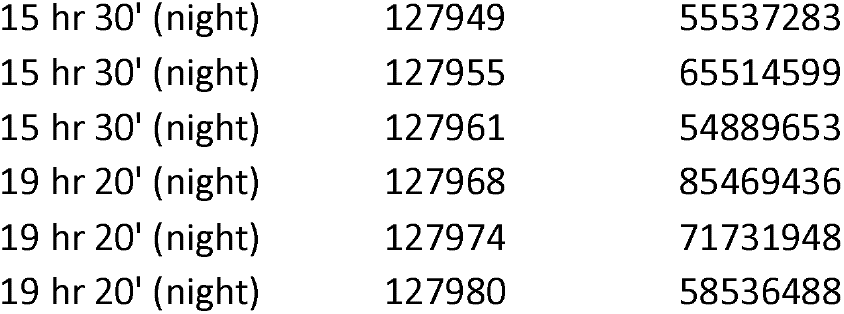
Detailed overview of the samples sequenced in this study.

| Condition | Age (days after infection) | Hours after artificial dawn | Replicate | Raw reads |
| --- | --- | --- | --- | --- |

|  |  |  | code |  |
| --- | --- | --- | --- | --- |
| Control | 14 | 5 hr 50' (day) | 127925 | 41193025 |
|  |  | 5 hr 50' (day) | 127935 | 50545664 |
|  |  | 5 hr 50' (day) | 127936 | 35205326 |
|  |  | 10 hr 40' (day) | 127906 | 56725609 |
|  |  | 10 hr 40' (day) | 127912 | 54273366 |
|  |  | 10 hr 40' (day) | 127918 | 38720139 |
|  |  | 15 hr 30' (night) | 127926 | 41309013 |
|  |  | 15 hr 30' (night) | 127931 | 55027843 |
|  |  | 15 hr 30' (night) | 127937 | 38004169 |
|  |  | 19 hr 20' (night) | 127907 | 39958547 |
|  |  | 19 hr 20' (night) | 127913 | 37101449 |
|  |  | 19 hr 20' (night) | 127919 | 40693049 |
| Infected | 14 | 5 hr 50' (day) | 127964 | 48508205 |
|  |  | 5 hr 50' (day) | 127970 | 41810661 |
|  |  | 5 hr 50' (day) | 127976 | 54739923 |
|  |  | 10 hr 40' (day) | 127946 | 42594195 |
|  |  | 10 hr 40' (day) | 127952 | 41155738 |
|  |  | 10 hr 40' (day) | 127958 | 58101982 |
|  |  | 15 hr 30' (night) | 127965 | 43247574 |
|  |  | 15 hr 30' (night) | 127971 | 46509846 |
|  |  | 15 hr 30' (night) | 127977 | 57602784 |
|  |  | 19 hr 20' (night) | 127947 | 40263406 |
|  |  | 19 hr 20' (night) | 127953 | 41398461 |
|  |  | 19 hr 20' (night) | 127959 | 50627500 |
| Control | 21 | 5 hr 50' (day) | 127908 | 52533389 |
|  |  | 5 hr 50' (day) | 127914 | 36118824 |
|  |  | 5 hr 50' (day) | 127941 | 64871718 |
|  |  | 10 hr 40' (day) | 127928 | 41179331 |
|  |  | 10 hr 40' (day) | 127933 | 45662748 |
|  |  | 10 hr 40' (day) | 127939 | 54002775 |
|  |  | 15 hr 30' (night) | 127909 | 43278660 |
|  |  | 15 hr 30' (night) | 127915 | 43201375 |
|  |  | 15 hr 30' (night) | 127920 | 42466381 |
|  |  | 19 hr 20' (night) | 127929 | 43287455 |
|  |  | 19 hr 20' (night) | 127934 | 46270564 |
|  |  | 19 hr 20' (night) | 127940 | 53657896 |
| Infected | 21 | 5 hr 50' (day) | 127948 | 59286277 |
|  |  | 5 hr 50' (day) | 127954 | 62740880 |
|  |  | 5 hr 50' (day) | 127960 | 46195940 |
|  |  | 10 hr 40' (day) | 127967 | 48074903 |
|  |  | 10 hr 40' (day) | 127973 | 61491275 |
|  |  | 10 hr 40' (day) | 127979 | 62580301 |

|  |  |  |
| --- | --- | --- |
| 15 hr 30' (night) | 127949 | 55537283 |
| 15 hr 30' (night) | 127955 | 65514599 |
| 15 hr 30' (night) | 127961 | 54889653 |
| 19 hr 20' (night) | 127968 | 85469436 |
| 19 hr 20' (night) | 127974 | 71731948 |
| 19 hr 20' (night) | 127980 | 58536488 |

**Additional File 3.**
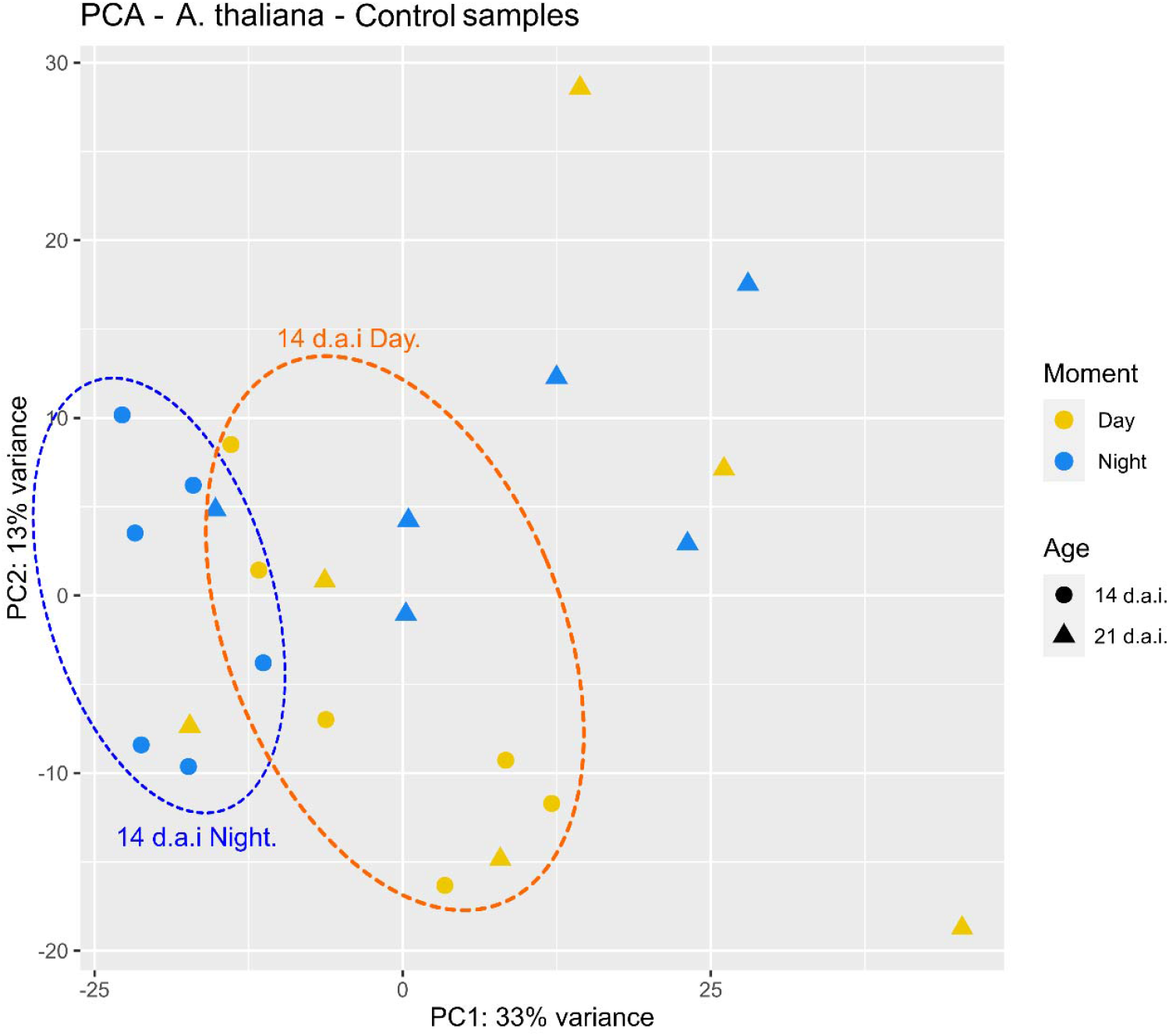
PCA on control A. thaliana plants, showing the decreased influence of the temporal factor (i.e., day and night) on the overall gene expression levels with the ageing of the plants.

**Additional File 4.**
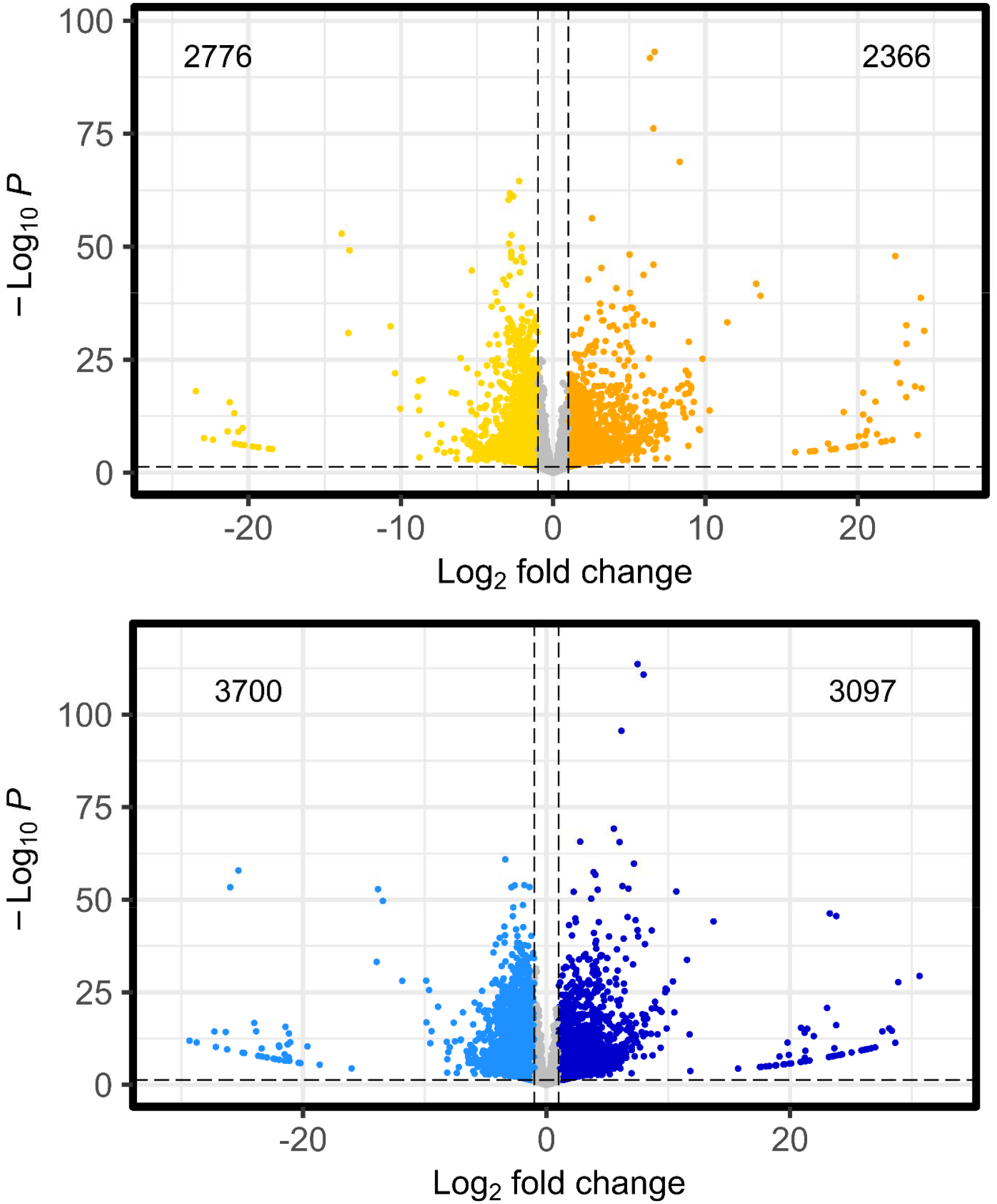
Volcano plot showing the number of DEGs in the day (top figure) and night (bottom figure) in infected A. thaliana plants 14 DAI

**Additional File 5.**
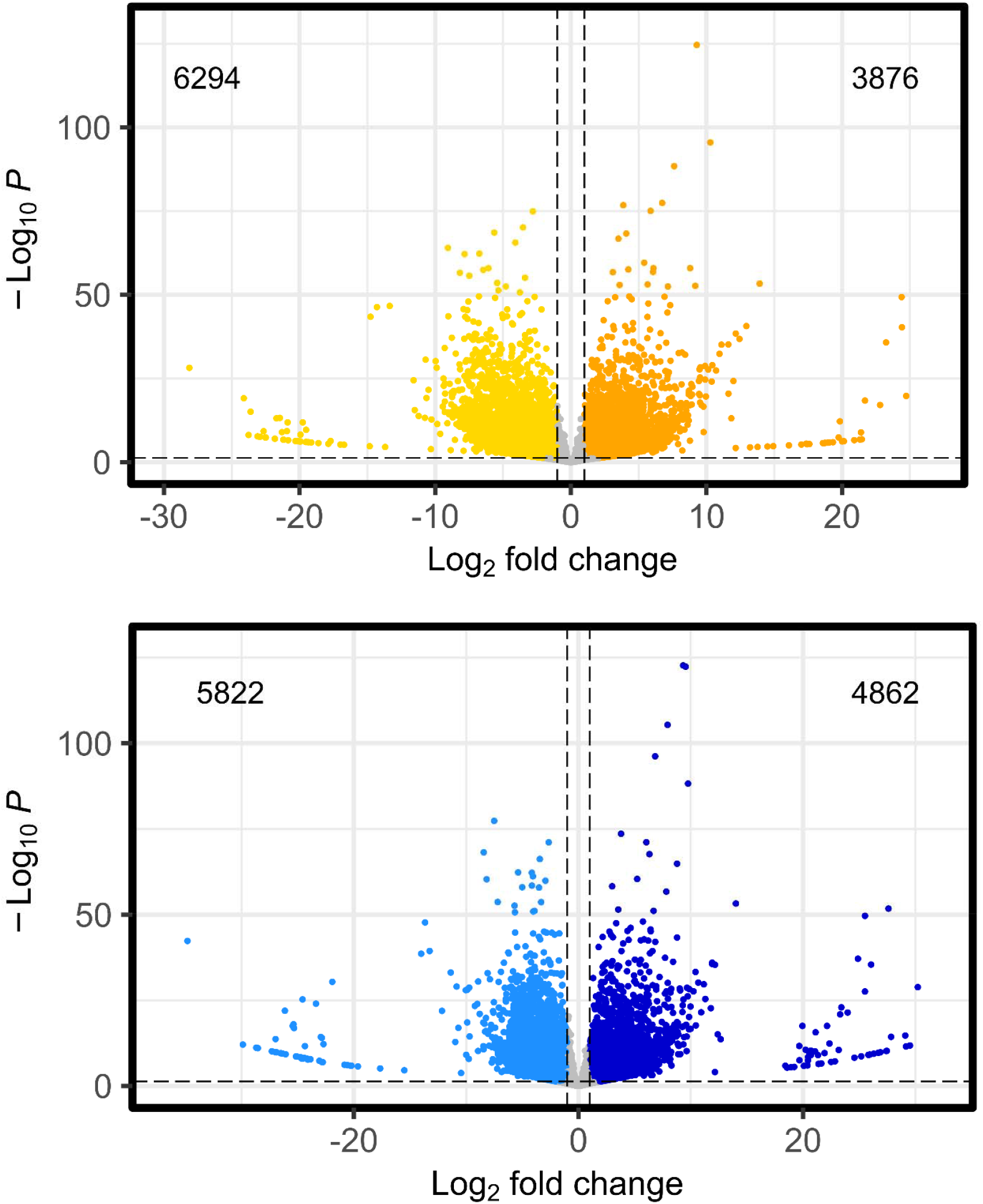
Volcano plot showing the number of DEGs in the day (top figure) and night (bottom figure) in infected A. thaliana plants 21 DAI

**Additional File 6.**
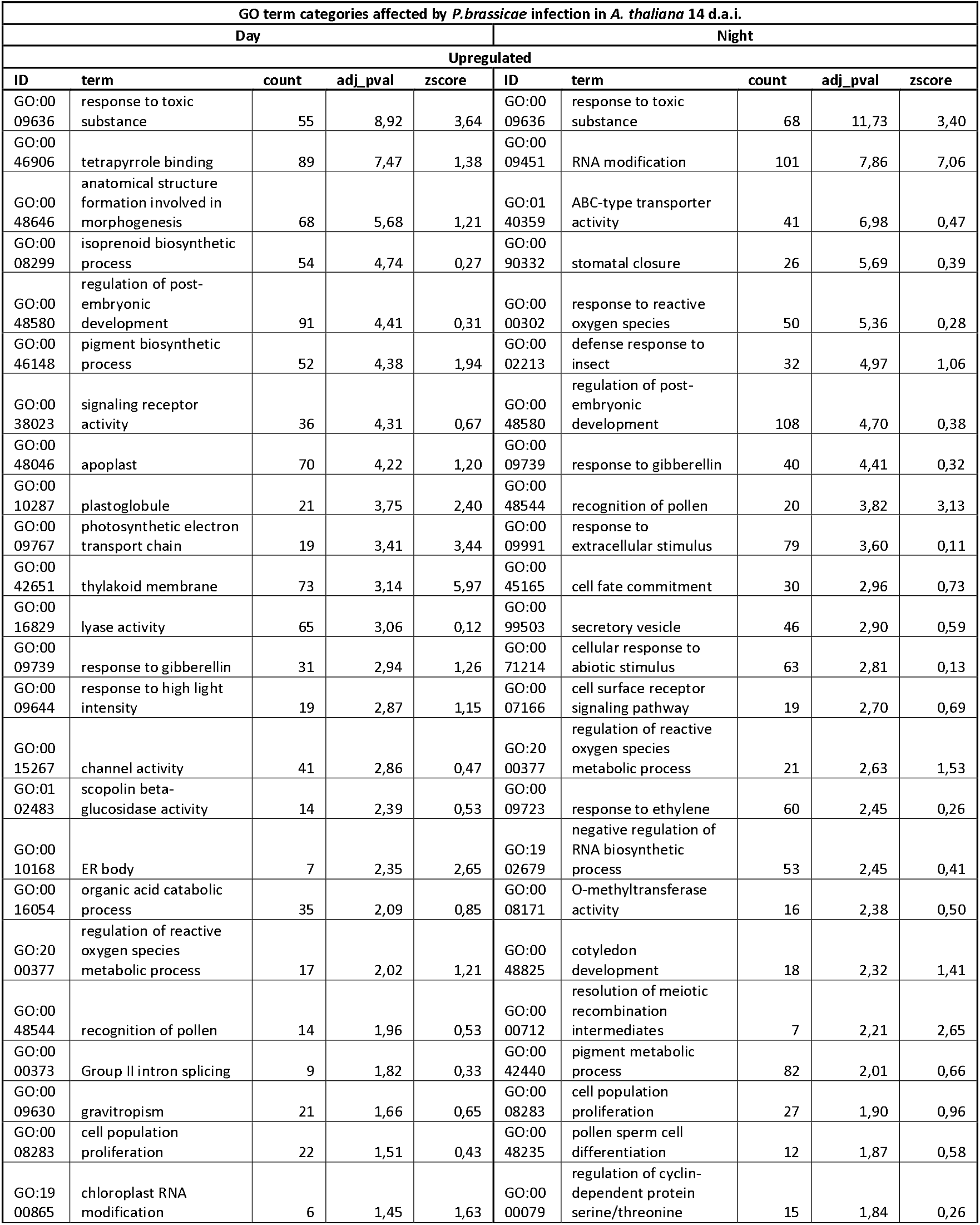
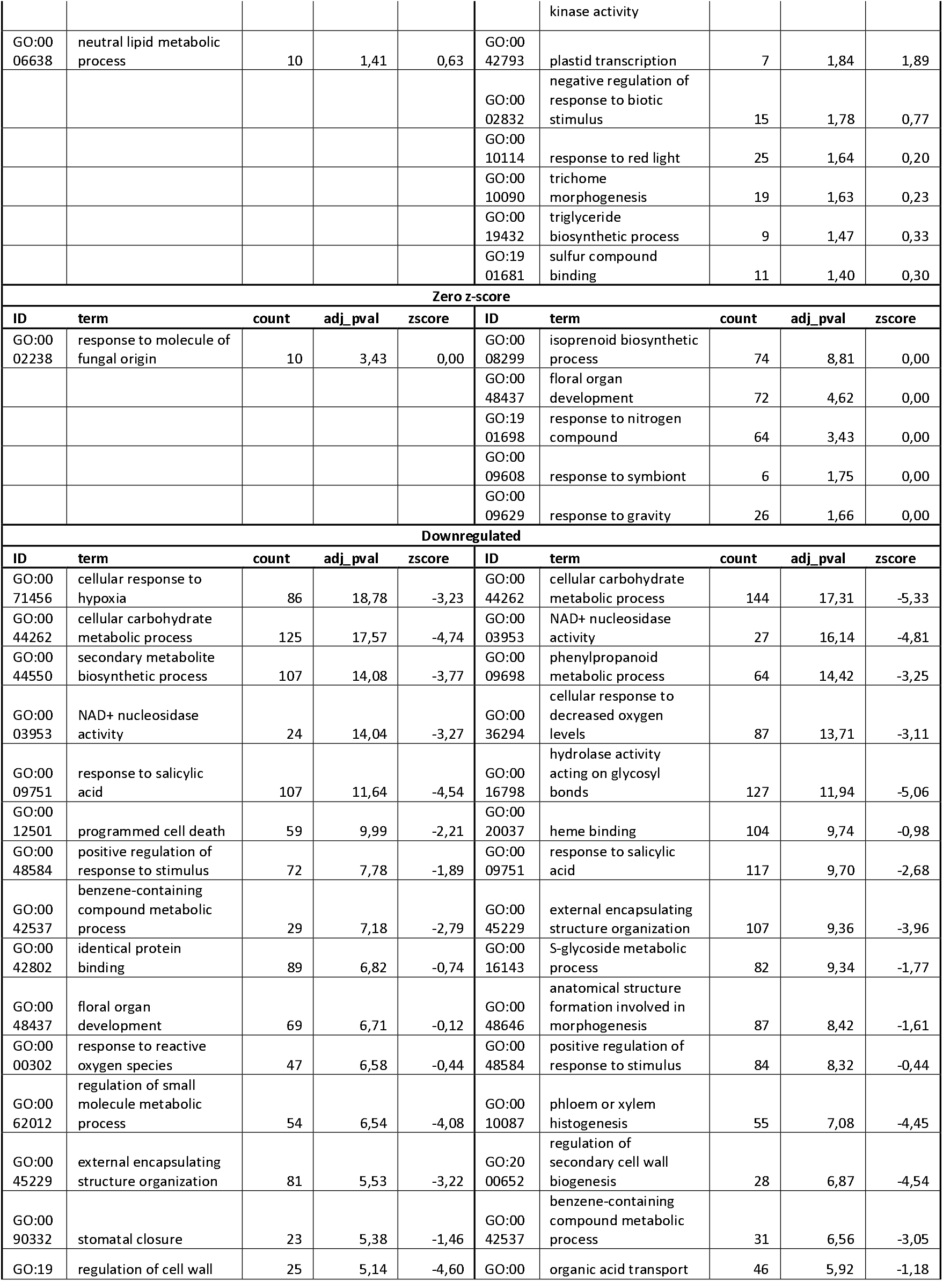
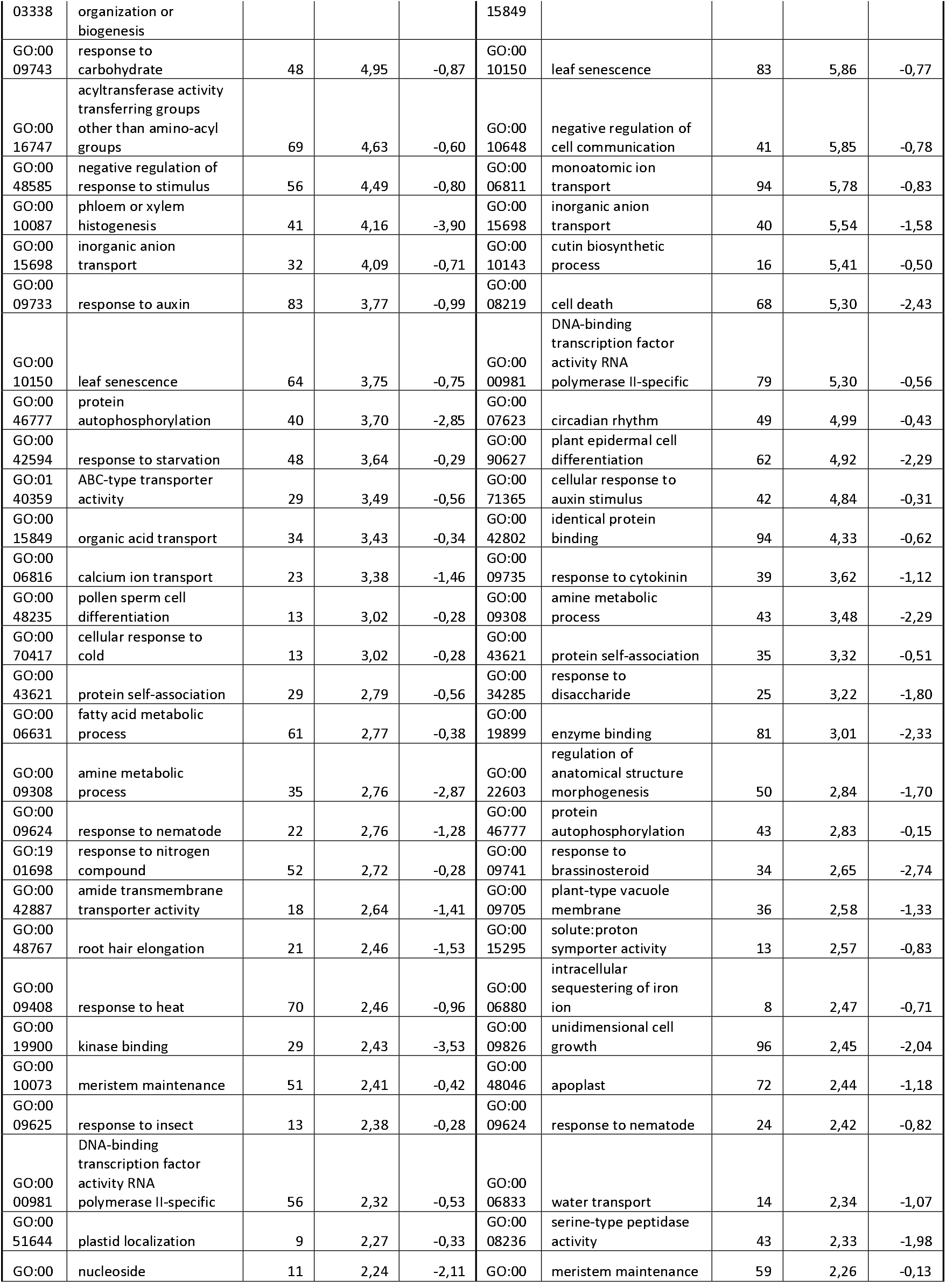
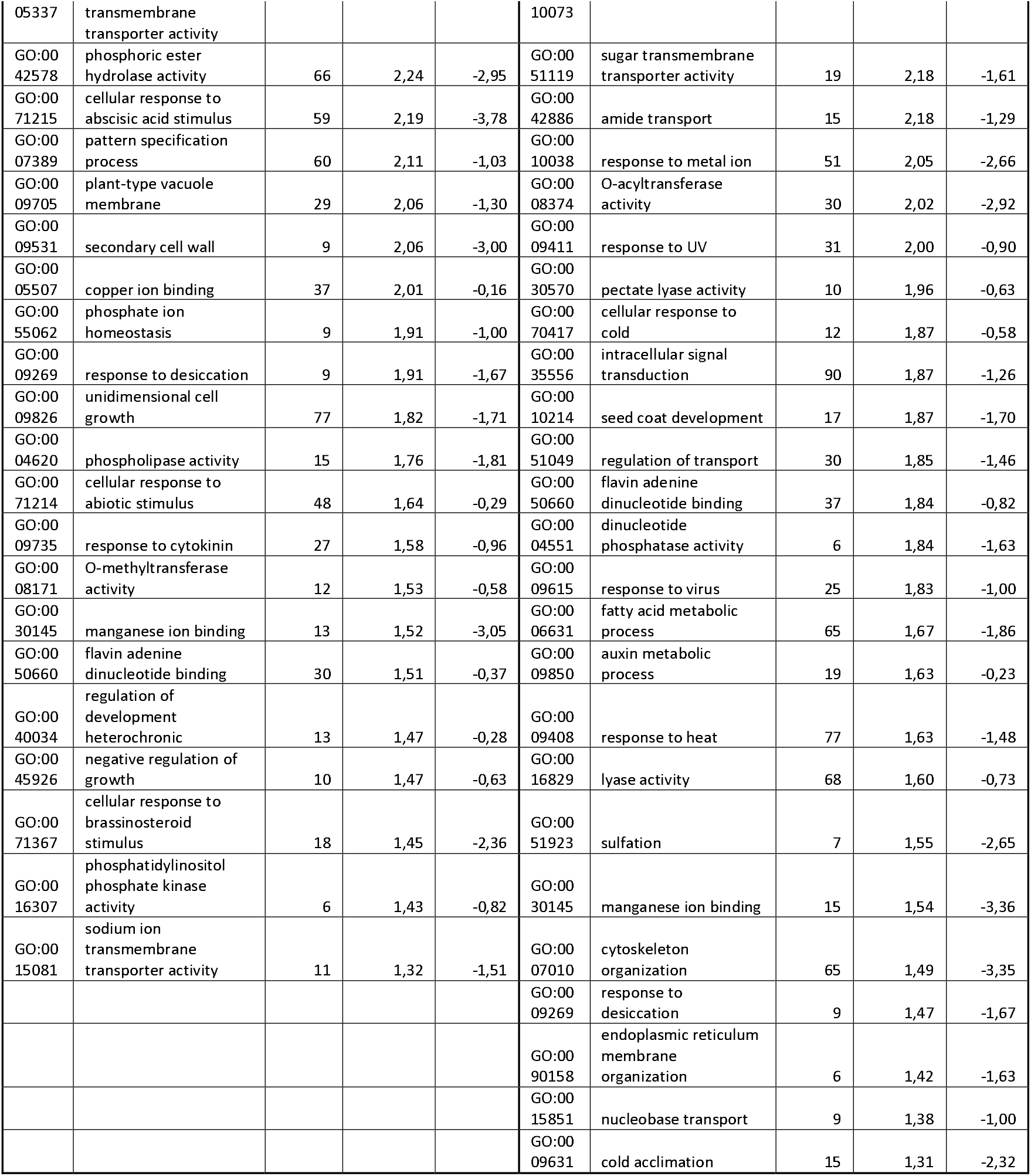
All GO categories enriched in DEGs in A. thaliana roots 14 DAI infected by P. brassicae as in Fig 3. GO categories are organised by decreasing -log10 adjusted p-value (i.e., top to bottom in the relative figure).

**Additional File 7.**
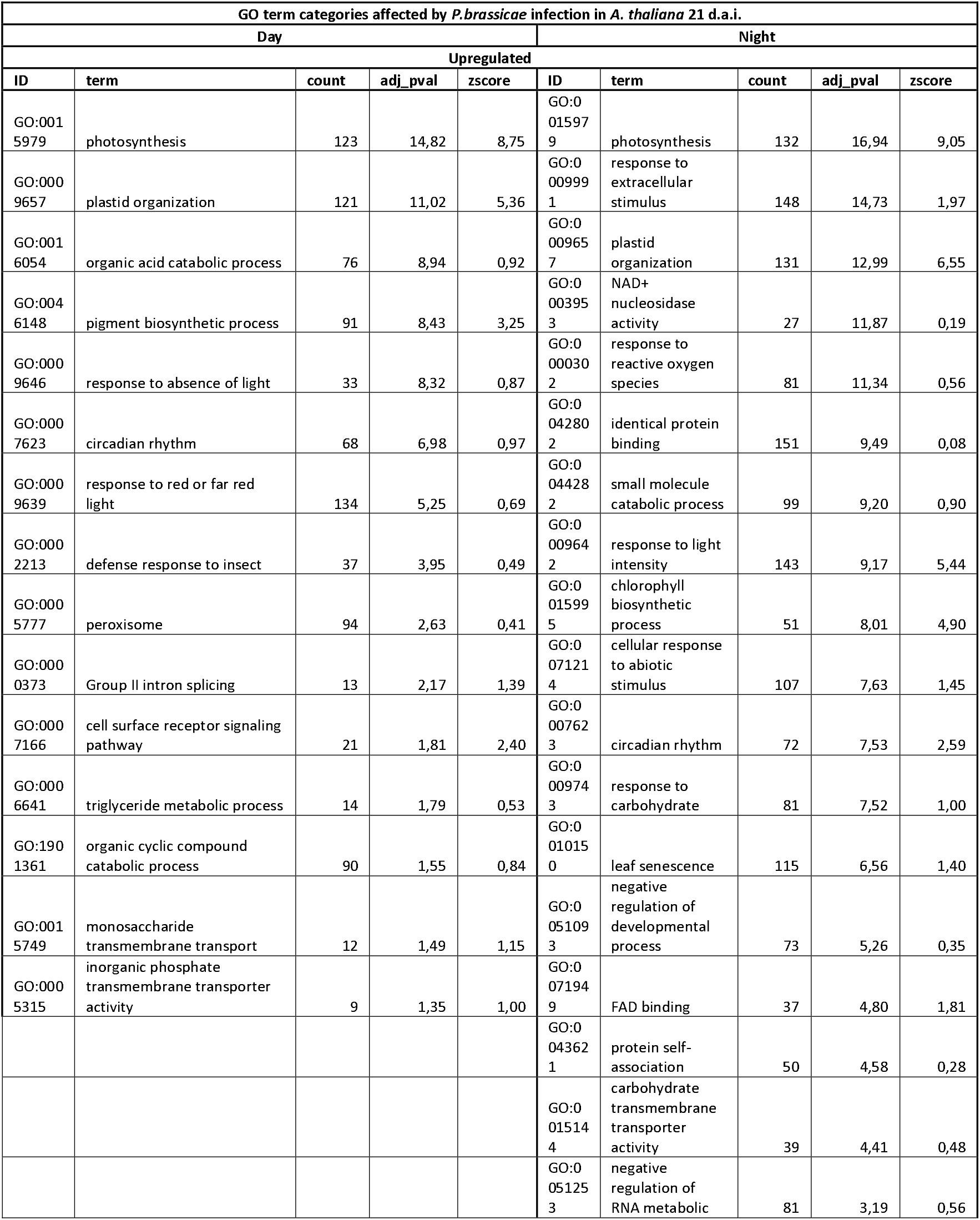
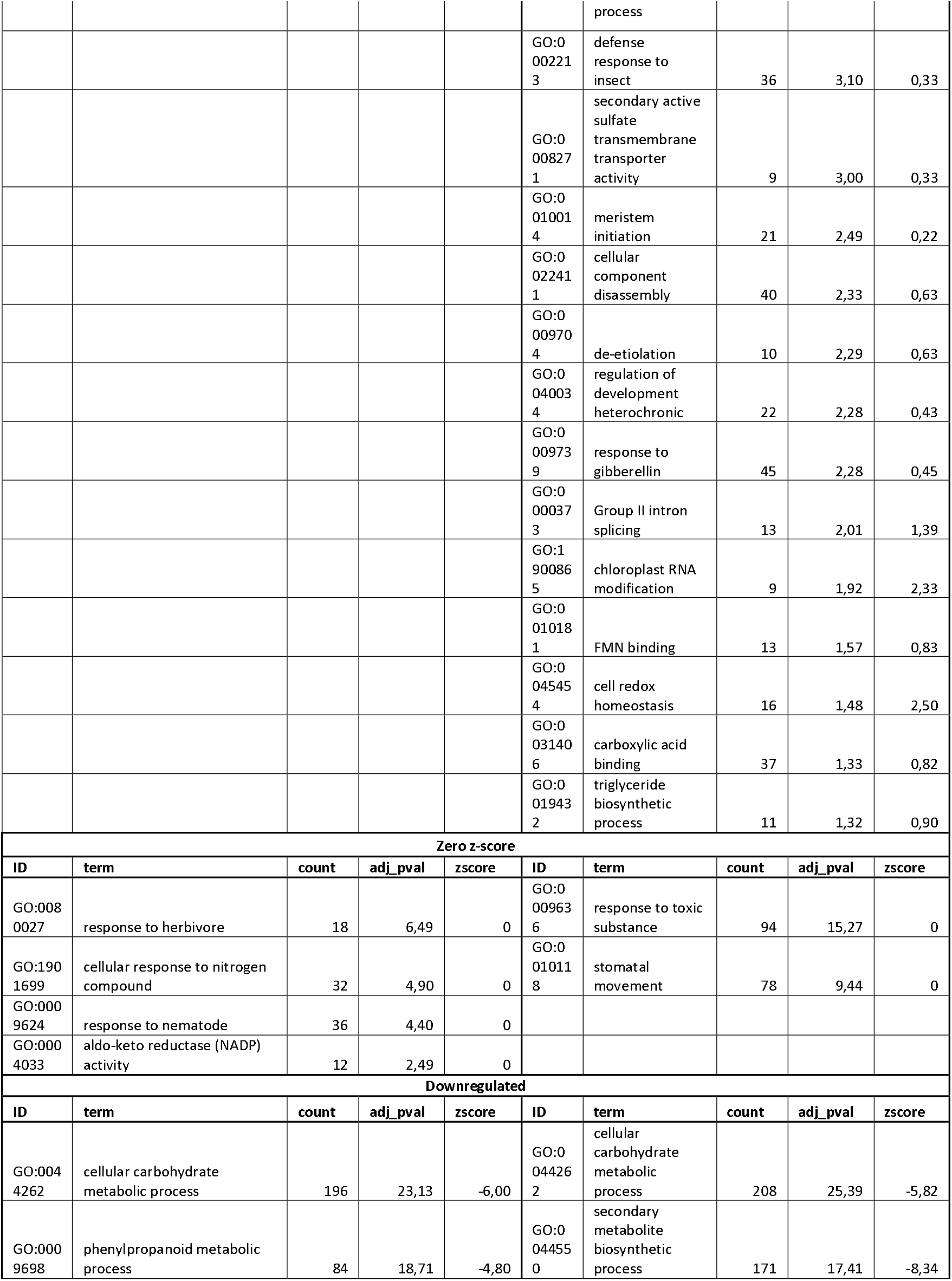
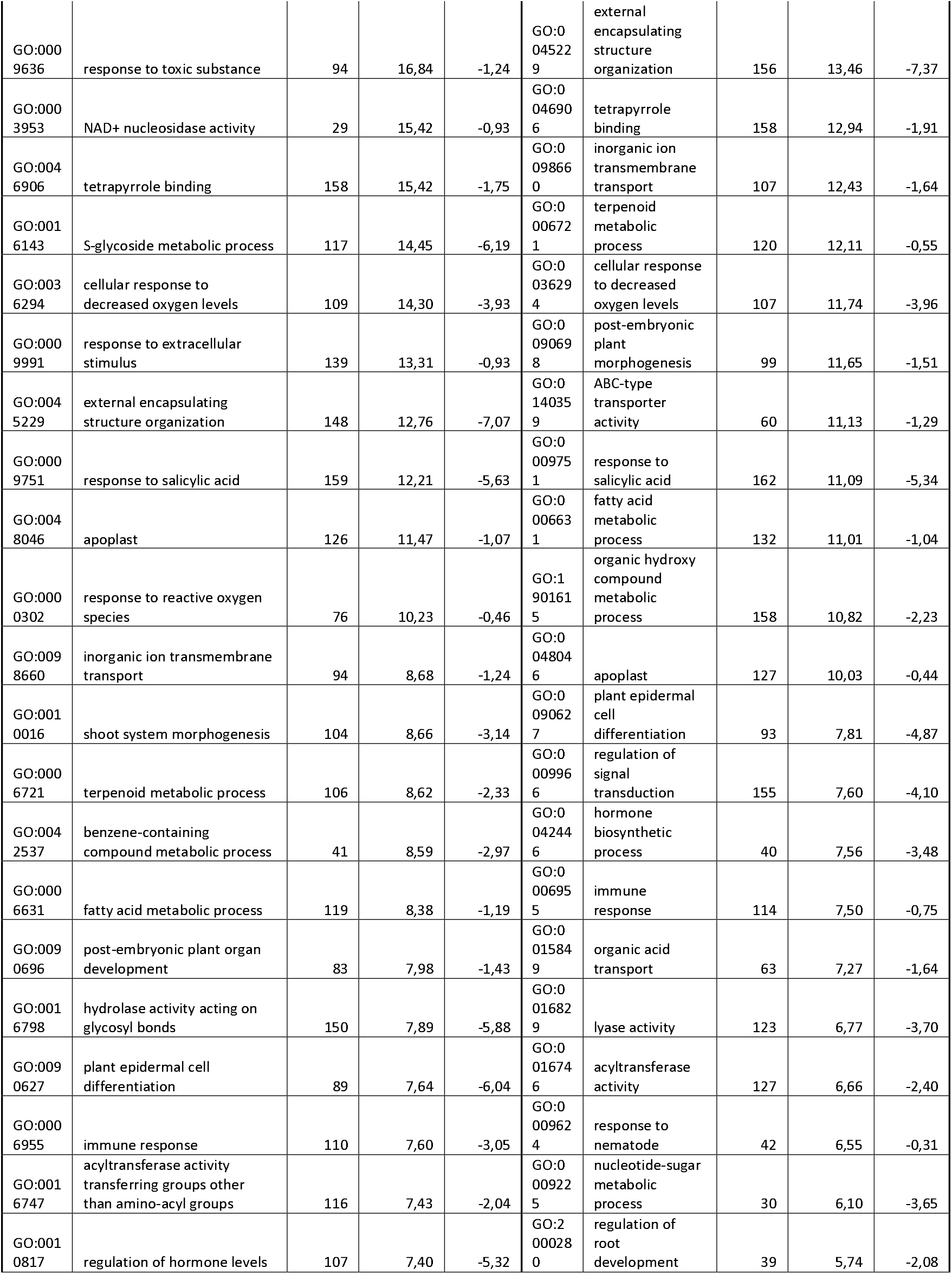
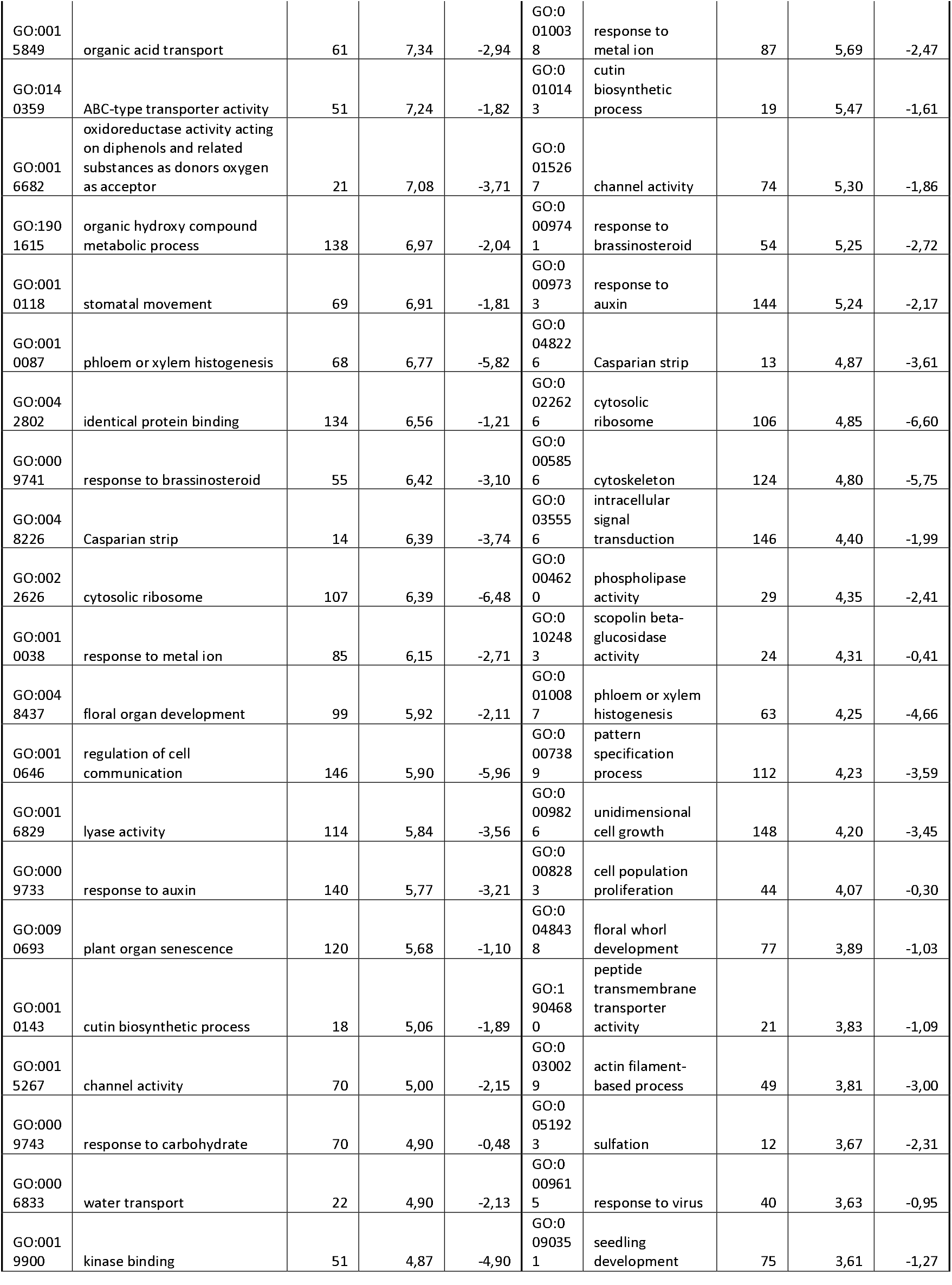
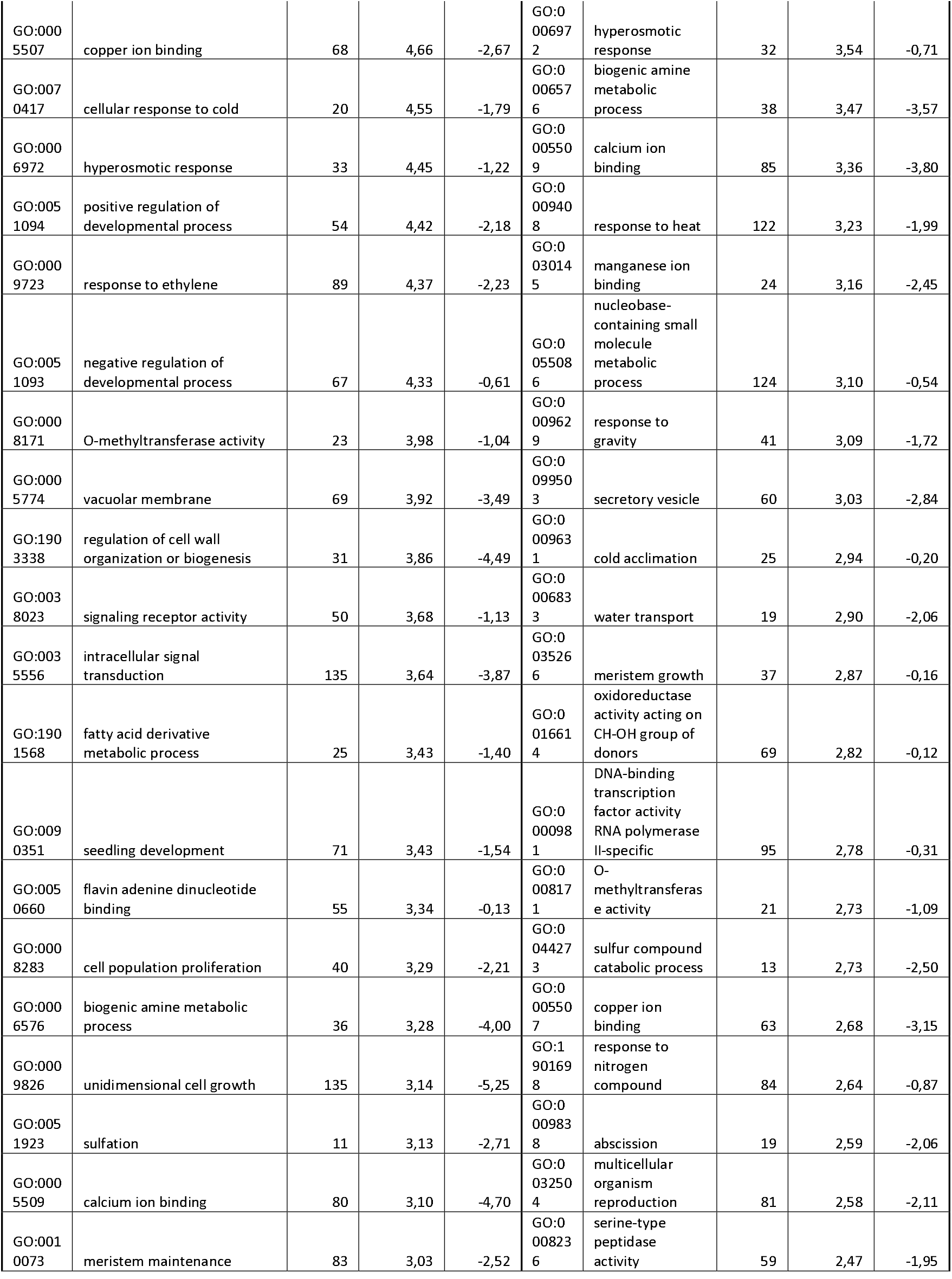
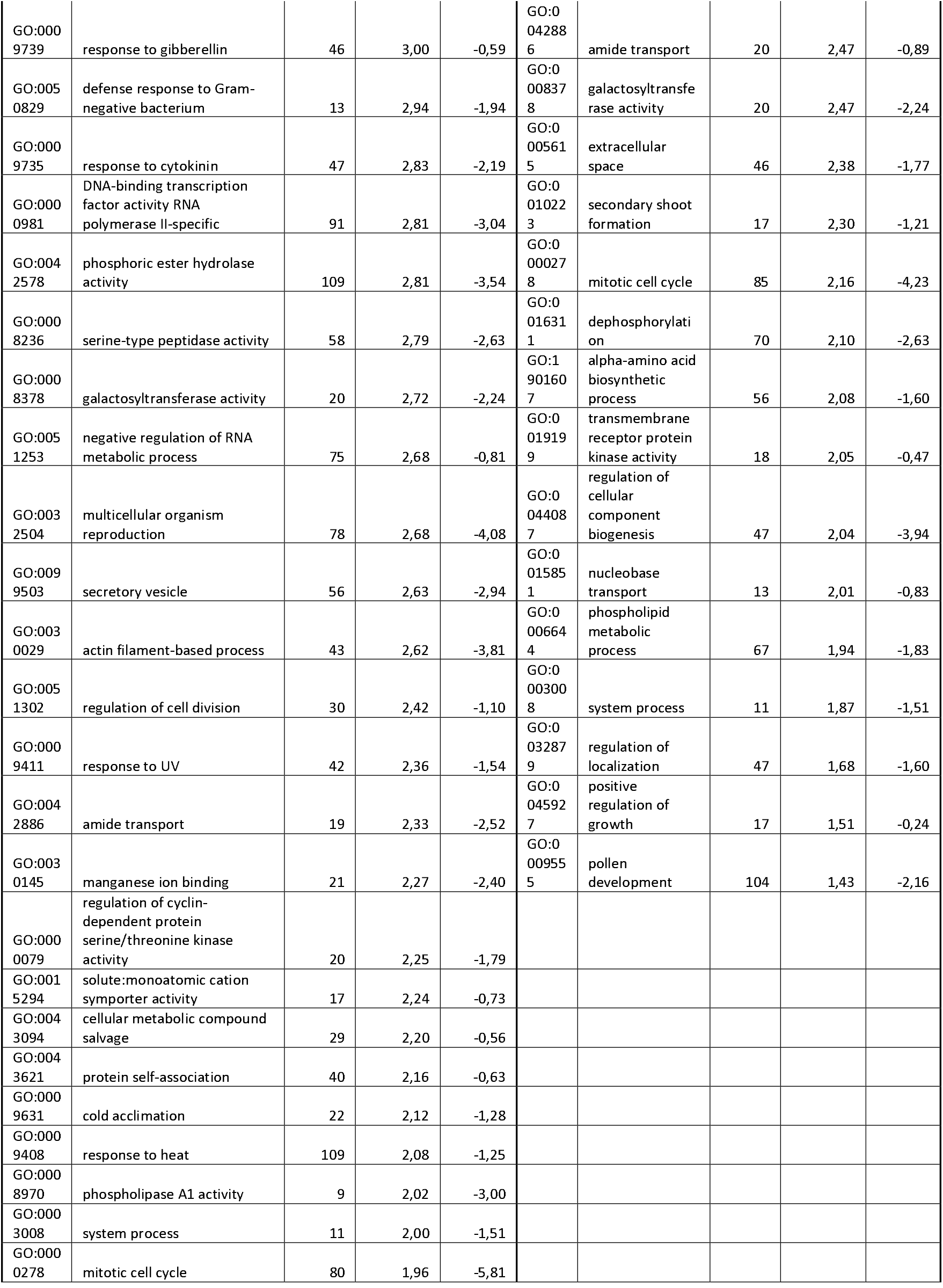
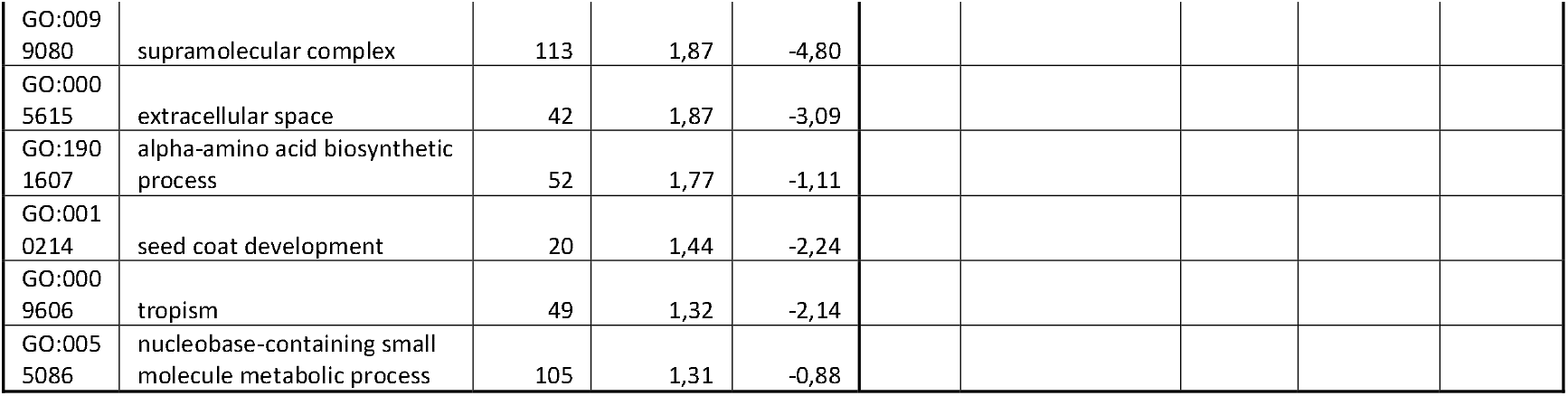
All GO categories enriched in DEGs in A. thaliana roots 21 DAI infected by P. brassicae as in Fig 3. GO categories are organised by decreasing -log10 adjusted p-value (i.e., top to bottom in the relative figure).

**Additional File 8.**
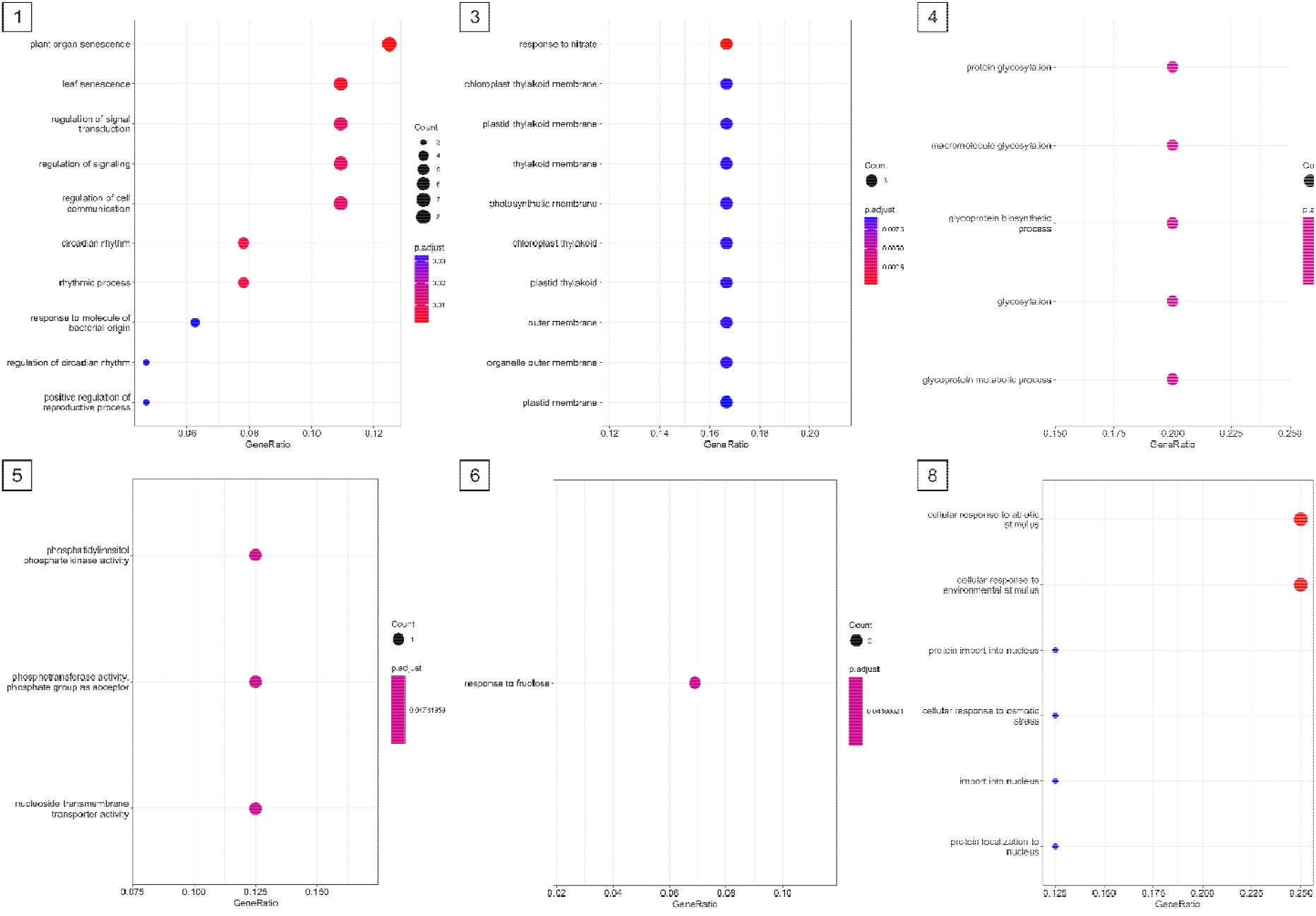
GO terms overrepresentation analysis of clusters of DEGs driven by the interaction between infection and time of the day 14 DAI

**Additional File 9.**
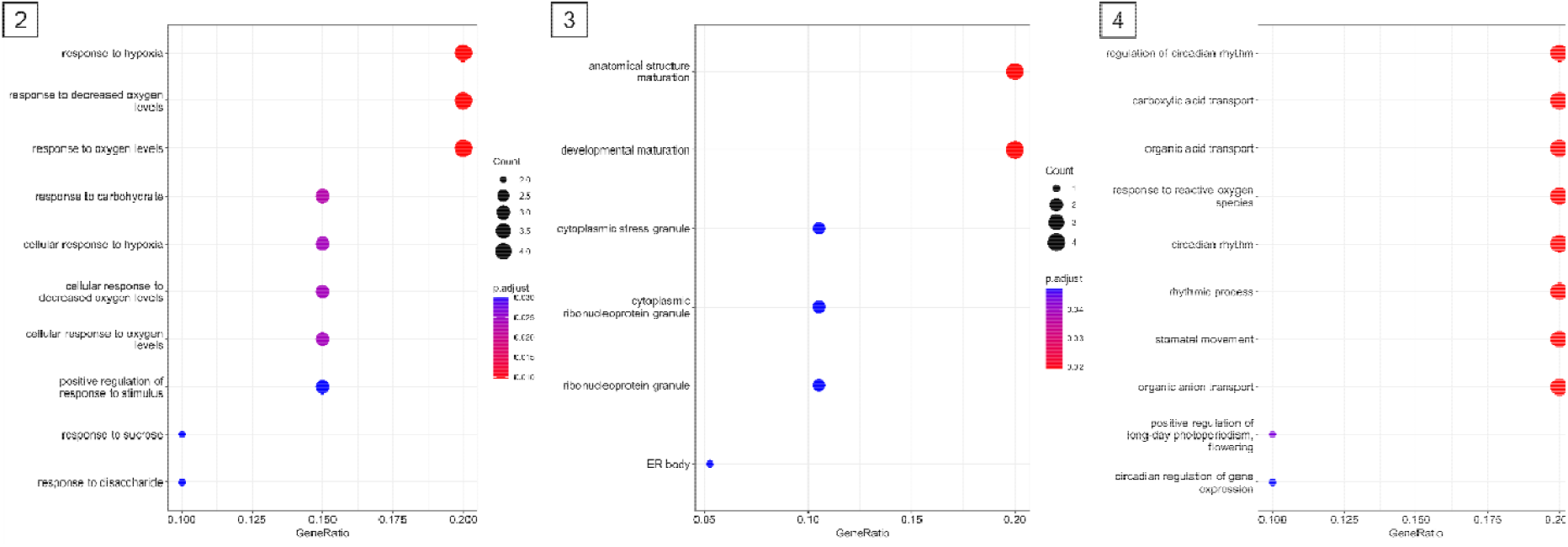
GO terms overrepresentation analysis of clusters of DEGs driven by the interaction between infection and time of the day 21 DAI

https://fileshare.uibk.ac.at/f/03d6ddfe1e6145bca1a6/?dl=1

Additional File 10 Enriched GO terms and corresponding genes underpinning figure 2 and 3 before data reduction.

